# High-throughput biodiversity surveying sheds new light on the brightest of insect taxa

**DOI:** 10.1101/2024.10.25.620209

**Authors:** Ela Iwaszkiewicz-Eggebrecht, Robert M. Goodsell, Bengt-Åke Bengsson, Marko Mutanen, Mårten Klinth, Laura van Dijk, Piotr Lukasik, Andreia Miraldo, Anders F. Andresson, Ayco J. M. Tack, Tomas Roslin, Fredrik Ronquist

## Abstract

Sampling of species-rich taxa followed by DNA metabarcoding is quickly becoming a popular high-throughput method for biodiversity inventories. Unfortunately, we know little about its accuracy and efficiency, as the results mostly pertain to poorly-known organism groups in underexplored environments or regions of the world. Here we ask what an extensive sampling effort based on Malaise trapping and metabarcoding can tell us about the lepidopteran fauna of Sweden – one of the best-understood insect taxa in one of the most-surveyed countries of the world. Specifically, we deployed 197 Malaise traps for a single year across Sweden in a systematic sampling design, then metabarcoded the resulting 4,749 bulk samples, and compared the results to existing data sources. We detected more than half (1,535) of the 2,990 lepidopteran species ever recorded as occurring in Sweden, and 323 species not reported during the sampling period by other data providers. Full-length barcoding of individual specimens confirmed three new species for the country and extensive range extensions for two species. It also corroborated eight genetically distinct COI variants that may represent new species to science, one of which has since been described. Most of the new records are for small and inconspicuous species and poorly surveyed regions, suggesting that they represent previously overlooked components of the fauna. Our findings, corroborated by independent metagenomic analyses, show that DNA metabarcoding can be a highly efficient and accurate method of biodiversity sampling, to the extent that it can generate significant new discoveries even for the most well-known of insect faunas.

## Introduction

Nature enthusiasts invest millions of hours and unparalleled know-how in generating biological records. Such data form the basis of national checklists, trend analyses, and ultimately of Red Lists guiding conservation priorities (Maes et al., 2015; Pescott et al., 2015). Nonetheless, current attention by naturalists comes with biases in space, time, and taxonomy (Rocha-Ortega et al., 2021). As a key concern, analyses of such opportunistically-recorded data are fraught with biases, as the underlying sampling effort will vary but is hard to characterize (Isaac & Pocock, 2015; Johnston et al., 2020). Larger, easier to spot, or more charismatic species receive a disproportionate amount of attention compared to highly diverse but inconspicuous taxa, such as most insect species. Recently, standardized sampling techniques coupled with DNA sequencing have proven efficient in revealing “dark diversity”, by shedding new light on taxa which are small, hyperdiverse and taxonomically neglected (Hartop et al., 2022; Srivathsan et al., 2023). To what extent the same methods may also brighten our view of the already well-lit parts of insect biodiversity remains poorly examined.

Of the methods available for accessing a comprehensive part of the local community, all have their drawbacks (Montgomery et al., 2021; van Klink et al., 2022, 2024). That a single method would sample *all* taxa with the same efficiency is an unrealistic expectation. In selecting a realistic solution, we need to account for at least two basic features of hyperdiverse taxa. First, given high diversity, we need to accumulate large numbers of individuals to characterize the many species in the community. Second, most species are rare, attesting to a need for intensive sampling in both space and time (Callaghan et al., 2023; Fisher et al., 1943; Goodsell et al., 2024). Both considerations call for the accumulation of large samples locally, and comprehensive coverage in space and time. As a workable compromise between these requirements, many recent projects have drawn on the Malaise trap (Malaise, 1937; Townes, 1972) as a wholesale approach to insect sampling (Karlsson, Hartop, et al., 2020; Li et al., 2023; Steinke et al., 2017).

With the sample in place, we need efficient means for taxonomic identification. Conventional identification by taxonomic experts using morphological characteristics is inappropriate for samples containing millions of specimens. As a vivid example, insect samples from a single tropical forest took about a month to generate, but a decade to identify (Basset et al., 2012). The sampling of around 1900 Swedish insect communities took three years to complete – but after two decades of intense work, less than 1% of the approximately 20 million specimens collected have been identified by expert taxonomists (Karlsson, Forshage, et al., 2020). As a key alternative to morphology-based identification, DNA metabarcoding enables the rapid identification of mass samples of insects. By extracting and sequencing a short fragment of DNA – the barcode – from a bulk insect sample and then comparing the DNA sequence to reference databases, we may achieve an efficient snapshot of the biodiversity present in a particular place at a particular time (deWaard et al., 2018). These metabarcoding methods are readily applicable to samples of thousands to millions of insects.

By leveraging advances in bioinformatics and molecular techniques, the coupling of mass-sampling of insects with high-throughput metabarcoding holds the potential to significantly enhance our ability to monitor and conserve insect biodiversity at a global scale (Hardulak et al., 2020; Ji et al., 2013). In fact, metabarcoding of environmental samples generated by Malaise samples is quickly becoming the go-to method for biodiversity inventories world-wide (deWaard et al., 2018; Li et al., 2024; Srivathsan et al., 2023). However, one of the biggest advantages of metabarcoding – that it can be applied to hyperdiverse and poorly known communities – is also its main limitation. As Malaise sampling and metabarcoding are routinely applied to samples and regions unamenable to other methods of identification, we rarely know the *right* answer in terms of what species should occur where, and what regional species pools we will be sampling. It is imperative that we test and scrutinize metabarcoding methods on well-known and comprehensively described taxa.

Lepidoptera, commonly known as butterflies and moths, provide a window of opportunity to validate the efficiency of mass sampling and metabarcoding methods, and to assess their overall efficiency in recovering the local species pool. Lepidoptera are among the best-known and described insect orders worldwide (https://gbif.org), and thus the expected composition of regional species pools is unusually well established. At the same time, Lepidoptera are sometimes assumed to be poorly sampled by Malaise traps (Busse et al., 2022; Karlsson, Forshage, et al., 2020; Matthews & Matthews, 1971) – thus making a Lepidoptera-based assessment a conservative test case for the efficiency of high-throughput surveying.

For a test region, Sweden provides the ideal setting – since the lepidopteran fauna of Sweden and neighboring countries is among the best-explored in the world (Ronquist et al., 2020; Roslin et al., 2022). Catalyzed by the activities of Linnaeus in the 18th century, the country has nurtured a long naturalist tradition, surviving until today and particularly noticeable in groups like Lepidoptera. In 2023, thousands of amateur and professional naturalists together reported an average of more than 120 occurrence records per species of Swedish Lepidoptera (https://gbif.org), at least an order of magnitude more than for any other group of insects. These naturalists sampled the national fauna by visual surveys and trapping methods, such as light traps, pheromone traps, and bait traps.

Despite this long-lasting effort, the inventory of Swedish Lepidoptera is hardly complete (Ronquist et al., 2020). New species are reported annually (Bengtsson, 2024; Palmqvist & Ryrholm, 2021, 2022) and old ones are split into multiple taxa (Aarvik et al., 2017).

There may still be undiscovered, resident species, as well as cryptic taxa within “known” taxa. Moreover, the fauna is far from static, and monitoring species’ ranges forms the basis of tracking regional change (Hällfors et al., 2024). The ultimate efficiency and coverage of the combined enthusiast effort remain unvalidated, since the actual ground truth is unknown. An alternative approach—if it were powerful—would offer a rare opportunity to calibrate the naturalists’ efforts and bring us closer to the full picture of lepidopteran biodiversity.

We deployed Malaise traps across Sweden for a single year (2019), leveraging metabarcoding techniques to uncover new taxa and expand our understanding of species distributions, in a project known as the Insect Biome Atlas (IBA; https://insectbiomeatlas.org). Here, we look into what such a systematic survey can teach us about the country’s lepidopteran fauna. Firstly, our inquiry explores what proportion of the known Swedish Lepidoptera fauna was covered by the IBA survey. Subsequently, we explore how the IBA data compare to the data collected by naturalists using traditional methods. Finally, we look into the new discoveries generated by IBA: Swedish Lepidoptera species recorded in locations beyond known distribution ranges, species new to Sweden, and putative species new to science.

## Materials and Methods

### Swedish species known to date

We retrieved the list of Lepidoptera records in the DynTaxa checklist (Backlund, 2024) from the web interface at https://namnochslaktskap.artfakta.se together with the occurrence status of each species, as identified by staff at the Swedish Species Information Centre (Artdatabanken; https://artdatabanken.se) (see data deposition info). DynTaxa (Backlund, 2024) lists all species that have been recorded from the country by professional and amateur naturalists—documented in natural history collections, in published papers, or in online databases—from the time of Linnaeus until the present.

### Occurrence data collected by naturalists

All major sources of data on Swedish Lepidoptera known to us are available through the Global Biodiversity Information Facility (GBIF; https://gbif.org). To represent the knowledge accumulated by traditional means up till the completion of the IBA survey, we downloaded all Swedish lepidopteran occurrence data from GBIF for the time period up until and including 2019. We restricted our download to human observations, preserved specimens and the Artportalen dataset (Species Observation System of Artdatabanken, https://artportalen.se) as it represents the bulk of occurrence data in GBIF.

### High-throughput biodiversity surveying

To generate records by high-throughput biodiversity surveying, we deployed Townes type Malaise traps at 197 sites spread throughout Sweden (IBA survey, Fig.1). Sites were selected in a systematic, hierarchical design, optimized to represent Swedish habitats and not to optimize the diversity or quantity of the insect catch. Traps were operated from January to December 2019 by citizen scientists, who replaced sample bottles weekly. This yielded a total of ∼4,749 Malaise trap samples (see Miraldo et al., 2024 for details).

**Fig. 1.**
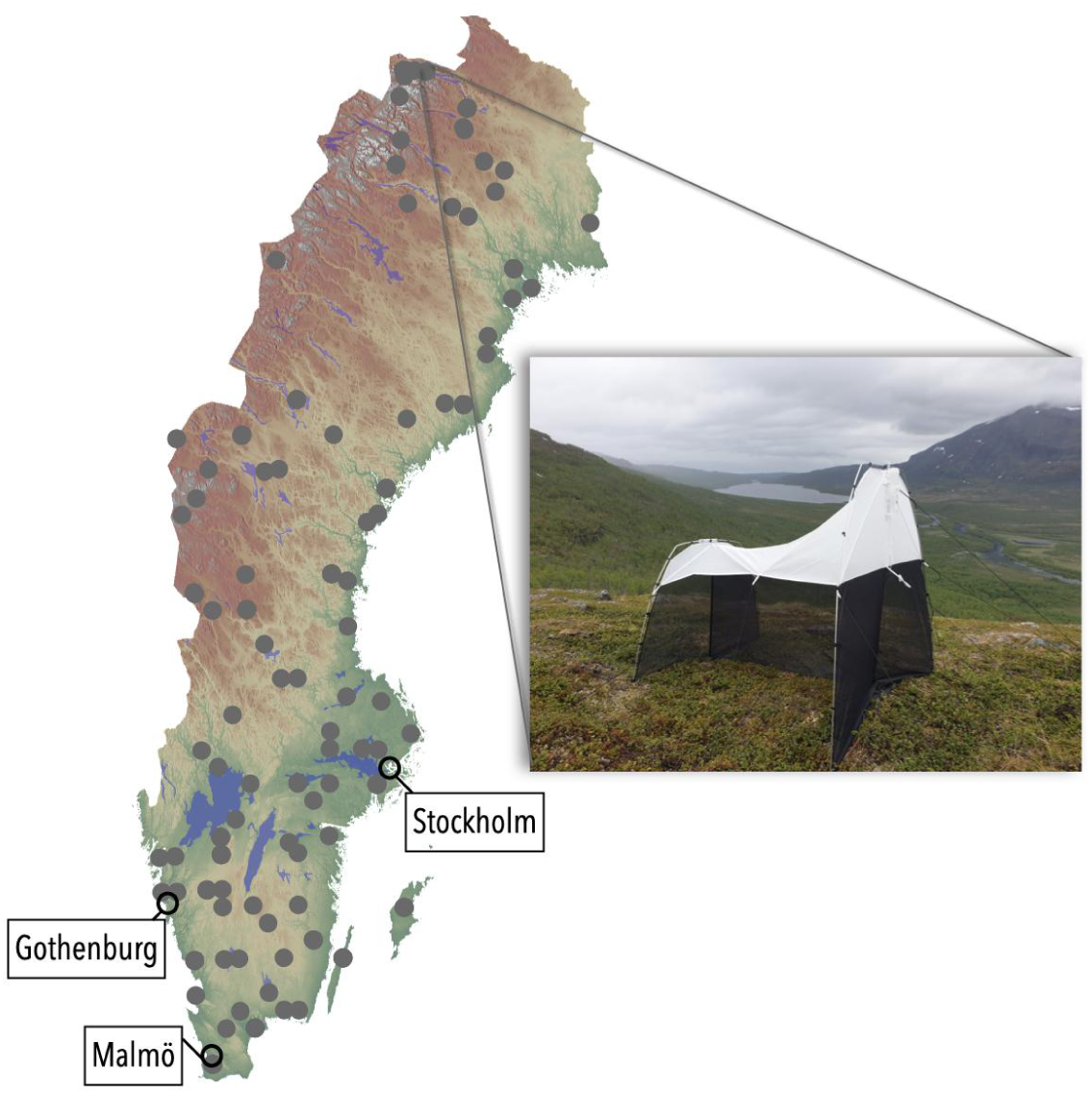
Distribution of IBA sampling sites in 2019 and main urban areas in Sweden with a photo showing one of the traps.

All samples were metabarcoded according to the FAVIS protocol (Iwaszkiewicz-Eggebrecht et al., 2023; Miraldo et al., 2024a). In short, we conducted a non-destructive mild lysis of the bulk sample, followed by DNA purification, preparation of amplicon libraries targeting a 418-bp fragment of the standard barcode region of the cytochrome *c* oxidase subunit I gene (COI), and sequencing the library pools on an Illumina NovaSeq 6000 platform using SPrime 500-cycle flow cells.

The resulting data was processed bioinformatically as described by Miraldo et al. (2024a). In short, after primer trimming, denoising, and chimera removal, all amplicon sequence variants (ASVs) were taxonomically annotated against a custom-made reference COI database (https://doi.org/10.17044/scilifelab.20514192.v4) using *SINTAX* (Edgar, 2016) and then clustered using *SWARM* (Mahé et al., 2014). Clusters were then quality-filtered by flagging noise (rare sequences) and likely nuclear DNA of mitochondrial origin (NUMTs) according to version 1 of the filtering algorithm described in Miraldo et al. (2024b) and available at [https://doi.org/10.17044/scilifelab.27202368.v1]. Finally, taxonomic annotations were matched to the list of Swedish species, and mismatches were analyzed in detail (see Supplementary Methods S1). We analyzed all clusters, including those flagged as potentially representing noise or NUMTs.

### Evaluating the efficiency of the high-throughput survey

We focused on three aspects: 1) *species coverage*, that is, the proportion of known Swedish Lepidoptera species recovered; 2) *comparison with naturalist data* with respect to taxonomic and regional coverage in the target year (2019); and 3) *new discoveries*, specifically the detection of putative new species, new records for Sweden, and records that significantly extended the known range of Swedish species. With respect to the third point, we note that the barcode reference libraries for Nordic Lepidoptera are nearly complete thanks to Finnish, Norwegian, German and North American efforts, among others (Roslin et al., 2022). Thus, lepidopteran clusters not matching existing reference libraries but passing all quality filtering represent potential new, previously undescribed species.

### Species coverage

We compared the list of species found in the IBA survey with the complete list of known Swedish lepidopteran species (DynTaxa) by matching species names. We did so for the entire fauna and at the family level. Here we sorted families by the traditional but phylogenetically unsupported split into those with mostly small species (“Microlepidoptera”) and those with mostly large species (“Macrolepidoptera”) (Wikipedia contributors, 2023b, 2023a, artfakta.se). To compare the taxonomic coverage of IBA survey and naturalist-collected data in one year, we filtered GBIF observations to include only records from 2019. Then we compared the number of occurrence records in the IBA data and in the GBIF dataset at the level of species and families.

### Comparing regional coverage of naturalist and IBA data in 2019

To compare the regional coverage of naturalist-collected and high-throughput survey data, we again used GBIF observations from 2019. We divided Sweden up into 125 roughly equally-sized polygons using *R* package *simple features* (Pebesma, 2018; Pebesma & Bivand, 2023) and then calculated the species richness of all Lepidopterans, Macrolepidopterans, and Microlepidopterans in each polygon for both naturalist-collected data and Malaise traps, whenever both data sources were available. We then calculated the proportion of the total occurrence records (i.e. the total number of unique species recorded in both datasets) recovered by each method to obtain a simple visual heuristic for comparison.

### Validation of new discoveries

To analyze the quality of the metabarcoding data in generating new discoveries, we selected 20 samples for validation with full-length barcoding of individual specimens from the original samples. The identification of the strongest cases for potential new species and new records for Sweden was based on a manual analysis of IBA clusters that did not match the DynTaxa checklist (Supplementary Results S1). To identify interesting new range records, we scanned the IBA data for species with at least one observation outside of their known range (according to GBIF data). For each lepidopteran species in GBIF, we calculated the observed range by computing the extent of occurrence (EOO) metrics via the minimum convex hull. We then calculated the longest distance between these ranges and the furthest record observed in the IBA data. We then selected 20 candidate species with the furthest distance from established EOO’s, double-checking GBIF ranges against historical literature records (Supplementary Material Methods S1). We made the final selection of bulk samples for specimen sorting and full-length barcoding using an *R* script combining criteria (minimal set of samples containing at least 50 reads of 20 candidate species representing all three categories: new species, new country records, range expansions).

### Individual specimen barcoding

The 20 bulk samples selected were sorted into 35 taxon fractions following the Swedish Malaise Trap Project protocols (Karlsson, Hartop, et al., 2020). We then searched the lepidopteran fraction for specimens matching the taxonomic identification of the candidate clusters. The selected specimens were processed with a rapid and efficient method of DNA barcoding described by Srivathsan et al. (2019). It combines non-destructive lysis, one-step PCR amplification with indexed primers and MinION sequencing followed by barcode selection using ONTbarcoder software (Srivathsan et al., 2021). For these validations, we targeted the full 658 bp “Folmer region” of the COI gene using LCO1490 and HCO2198 primers (Folmer et al., 1994) with unique indices attached (Srivathsan et al., 2021). The resulting barcode sequences were then matched against the BOLD database (Ratnasingham & Hebert, 2007) and BLASTed against GenBank data (Sayers et al., 2024).

## Results

The IBA high-throughput survey resulted in 544,263 amplicon sequence variants (ASVs) grouped into 33,888 clusters (proxy for species), 1,705 of which were annotated to Lepidoptera. The clustering of Lepidoptera sequences was highly consistent with species identifications based on traditional taxonomy; the errors, when they occurred, tended to be in the direction of splitting rather than lumping (precision 0.997, recall 0.980, homogeneity 0.997, completeness 0.991 after noise filtering; Sundh et al. 2024). Out of the 1,705 Lepidoptera clusters, 1,437 were successfully assigned to species using the default pipeline. This number increased to 1,636 after the manual resolution of problems with taxonomic name conflicts, and data quality problems in the BOLD reference library (Supplementary Methods S1).

### Species coverage

The known Swedish Lepidoptera fauna comprises 2,990 species belonging to 76 families; 2,697 of those are resident and regularly reproducing in the country (later referred to as *natural residents*). Some of the most species-rich families include the Tortricidae (leafroller moths) with over 400 species, Noctuidae (including Owlet moths) with around 350 species and Geometridae (including the Emeralds and Carpet moths) represented by about 320 species.

The IBA clusters matched 1,535 species known from Sweden. Thus, a single year of high-throughput biodiversity survey recovered 51% of all lepidopteran species known from the country (Fig. 2A). All but three species were natural residents; the survey thus covered 1,532 (57%) of the natural residents in the fauna.

**Fig. 2.**
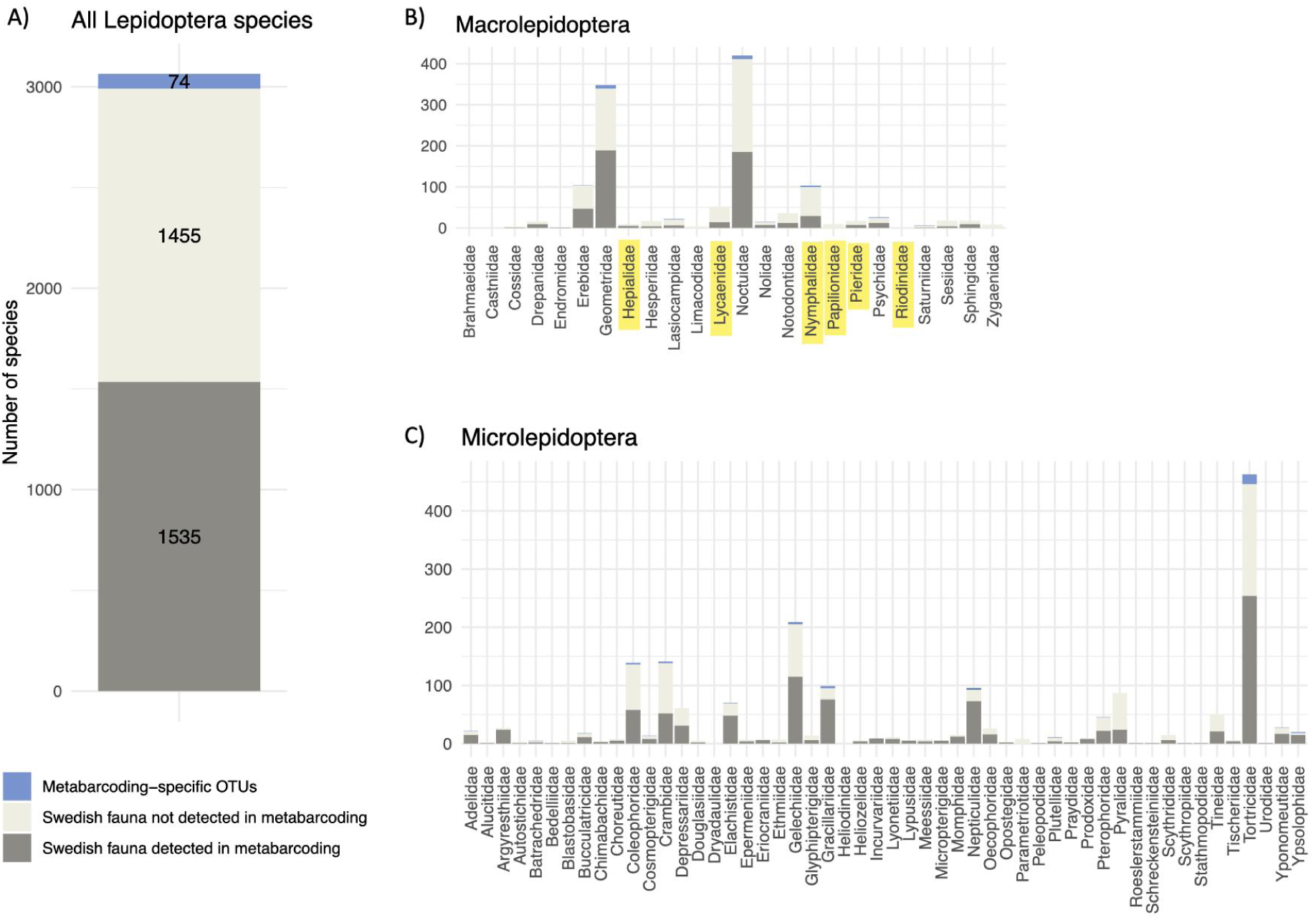
The proportion of the Swedish lepidopteran fauna detected by the high-throughput IBA survey using DNA metabarcoding of Malaise trap samples. A) across all taxa, B) per macrolepidopteran family and C) per microlepidopteran family. The dark gray sections of the bars correspond to Swedish species detected by the IBA survey; the light gray sections show species recorded from Sweden but not detected by IBA; and the blue sections show clusters (putative species) detected uniquely by IBA. Family names of butterflies are highlighted in yellow.

The proportion of taxa recorded was slightly lower across families of Macrolepidoptera when compared to Microlepidoptera (44% vs. 56%, Table S1) and tended to be even lower for butterflies (25%, Table S1; Fig. 2B and 2C, with butterfly families highlighted in yellow). Only nine of the 76 families were missing completely, all of them species-poor. The missing species are associated with significantly fewer GBIF records than the ones recovered in the IBA survey (two-sided t-test, p<0.001), indicating that they are on average less common (Supplementary Fig. S1).

### Comparison to data from naturalists

In 2019, naturalists reported 444,078 occurrence records of insects from Sweden; the Lepidoptera stood for 61% of those (272,380 records; Supplementary Fig. S2). The IBA data, viewed as species-space-time points with each point denoting the presence of a single cluster in a single sample, comprised 42,885 records, that is, 16% of the naturalist data for Lepidoptera. Lepidoptera was the only major insect group for which the naturalist data substantially exceeded that of the IBA survey viewed as simple occurrence records (Supplementary Fig. S2). The naturalist data from 2019 covered 2,379 species of Lepidoptera, 1,228 of which (52%) were also among the 1,535 known species recovered in the IBA data (Supplementary Fig. S3).

Taxonomically, the naturalist data are strongly biased towards families of butterflies and other macrolepidopterans (Fig. 3A). The amount of IBA occurrence data equals or exceeds the naturalist data (in terms of the average number of records per known species) for around half of the families of Microlepidoptera (Fig. 3A) and for many individual species of Microlepidoptera (Fig. 3B). At the species level, there is very weak correlation between the number of naturalist and high-throughput survey occurrence records (Pearson correlation coefficient (log scale) 0.11; Fig. 3B).

**Fig. 3.**
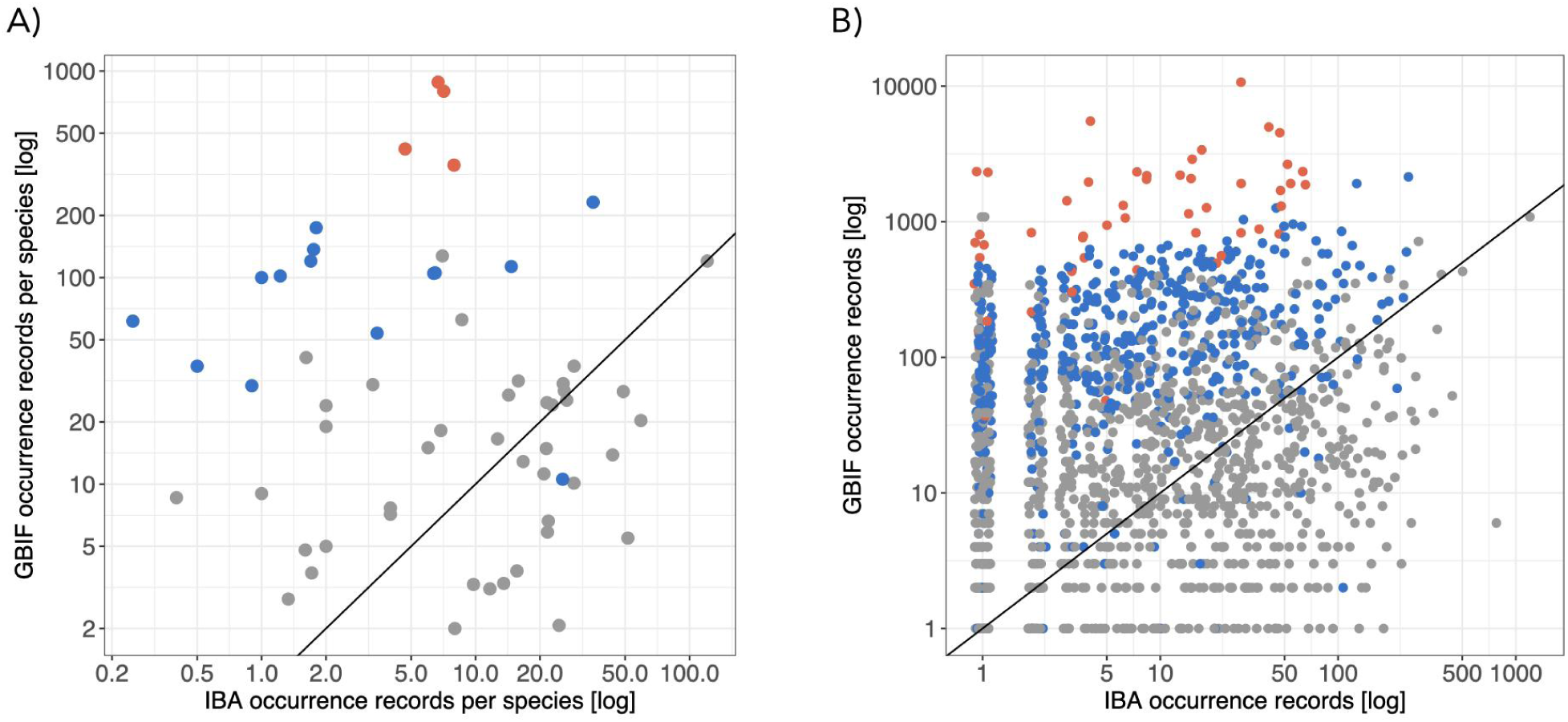
Comparison of the taxonomic coverage of naturalist (GBIF) and high-throughput survey (IBA) occurrence data in 2019. A) Average number of occurrence records per species in each family; B) Number of occurrence records for each species (with jitter to improve visualization). Butterflies in red, other macrolepidopterans in blue, microlepidopterans in grey. For points above the line, there is more occurrence data from naturalists; below the line, there is more data from the high-throughput survey.

Spatially, the amount of naturalist data largely appears to reflect human population density, with Stockholm, Mälardalen (lake area west of Stockholm), Gothenburg and Skåne (southern Sweden) standing out (Figs 4A and S4A). The naturalists excel at covering the species pools in these areas, particularly for butterflies and other Macrolepidoptera (Fig. 4B). The IBA data performed better than naturalists in recovering the local species pool in sparsely populated areas of the country. This is particularly noticeable for the Microlepidoptera (Fig. 4C).

**Fig. 4.**
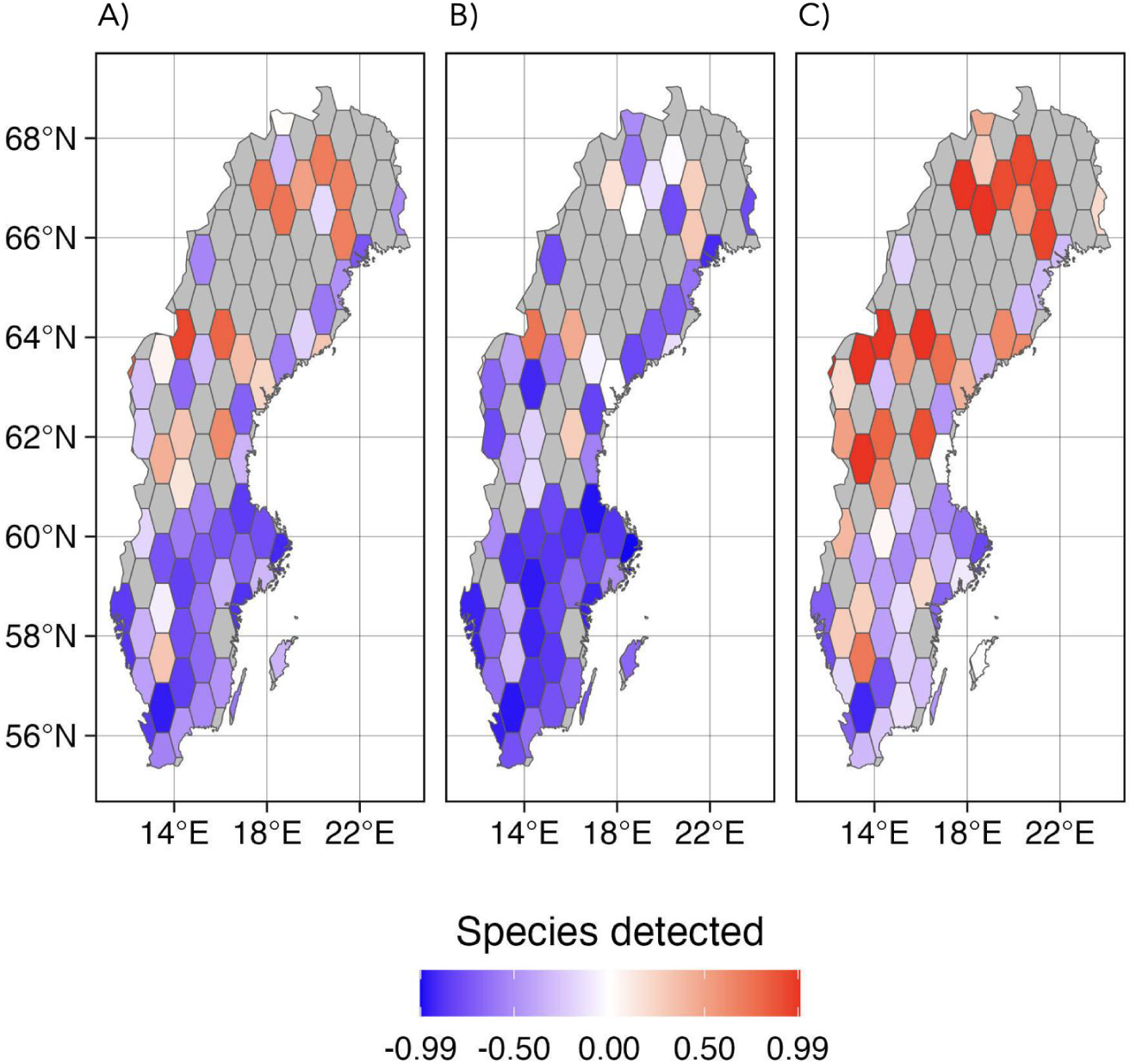
Comparison of the regional coverage of naturalist and high-throughput survey (IBA) data in 2019. A) Difference in the proportions of species coverage in each spatial bin across all groups. B) Ditto for macrolepidopteran fauna. C) Ditto for microlepidopteran fauna. The colors reflect the difference in the proportion of species retrieved by the two surveys within each polygon. Blue colors represent more observations retrieved from naturalists (GBIF data) and red colors more data from the IBA survey. Gray polygons are areas not covered by the IBA survey.

### Validation of new discoveries

The IBA data included 74 clusters that did not match any known Swedish species (Fig. 2A). Many of these belonged to small moth families (Microlepidoptera) but there were also potentially new clusters of Macrolepidoptera. In addition, the IBA data included several records that extended known species ranges considerably. Finally, there were 100 IBA clusters of Lepidoptera flagged by our bioinformatics pipeline as potential NUMTs or other noise.

After careful revision we selected 20 clusters representing such new or surprising records for validation with full-length barcoding. We examined 12 clusters that did not match records in reference libraries, and thus were potentially new to science (including six which had been flagged as noise). We also looked at four species not found in Sweden previously and four examples of conspicuous range expansions.

We were able to validate three of the four species new to Sweden, and two of the four range expansions—a third range expansion was also validated but turned out incorrect because our pipeline lumped two species and resolved the annotation as the more common southern species. Examples of the validated new discoveries are shown in Figure 5 (see Supplementary Results S1 and Table S2 for details).

**Fig. 5.**
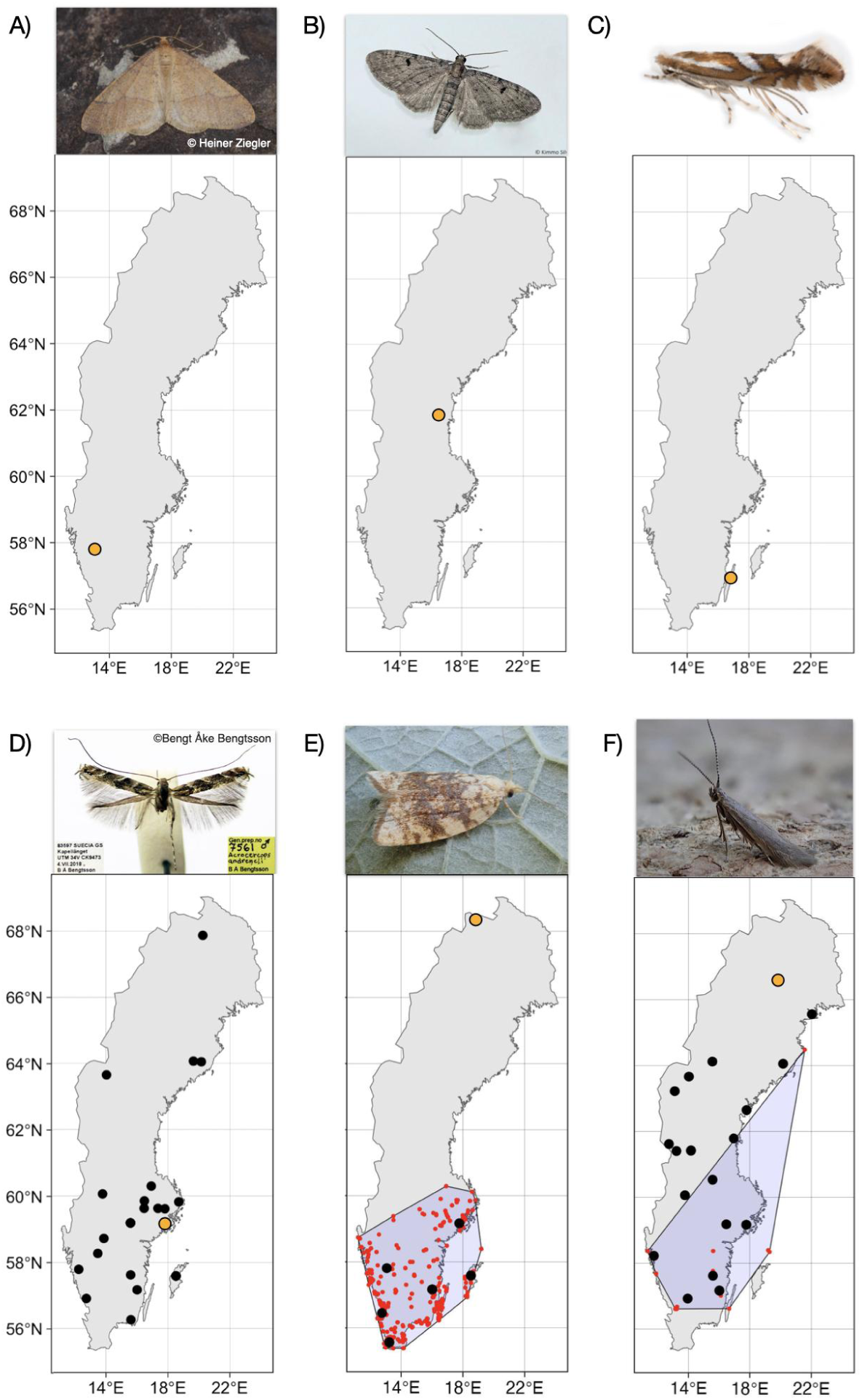
Examples of new discoveries resulting from the high-throughput IBA survey. Black dots indicate IBA occurrence records and orange dots mark IBA samples that were sorted and individually barcoded. A) *Agriopis budashkini* (Geometridae), a species described in 2009 from southeastern Europe, new to Sweden; B) *Eupithecia groenblomi* (Geometridae) and C) *Phyllonorycter mespilella* (Gracillariidae), both new to Sweden but known from neighboring countries; D) *Acrocercops andreneli* (Gracillariidae), a putative invasive species described in 2023; E) *Aleimma loeflingiana* (Tortricidae) and F) *Coleophora juncicolella* (Coleophoridae), both with known distribution significantly extended (previous records appear in red, surrounded by the minimal convex hull). Photo credits: A – © Heiner Ziegler – <u>some</u> <u>rights reserved (CC BY-NC)</u>, B – © Kimmo Silvonen https://www.suomen-perhoset.fi accessed 15/08/2024, C – © Chris Lewis https://britishlepidoptera.weebly.com accessed 22/10/2024, D – Bengt-Åke Bengtsson, E and F – Phil Barden https://www.ukmoths.org.uk accessed 15/08/2024.

Eight clusters putatively representing new species, including three of six clusters flagged as noise and five of six flagged as authentic COI sequences, matched barcodes that were generated for individual specimens sorted out from Malaise traps (Supplementary Table S2). The eight validated clusters represent genetically distinct COI variants, separated from known variants at distances suggesting they come from separate species. One of these variants apparently corresponds to *Acrocercops andreneli*, an invasive species that was not described until last year (Nel et al., 2023) It was first reported from Sweden in 2024 (Bengtsson, 2024), and the IBA data represent a unique record of its distribution in the country in 2019 (Fig. 5D).

## Discussion

Malaise traps are frequently assumed to be poorly-suited for sampling Lepidoptera and DNA metabarcoding is dismissed as a surveying method due to the supposed poor data quality (Busse et al., 2022; Karlsson, Hartop, et al., 2020; Matthews & Matthews, 1971). However, other findings challenge these notions (Aagaard et al., 2017; Rosa et al., 2019; Takano et al., 2024). Similarly, an analysis of the Swedish Insect Inventory Project (SIIP) material using traditional morphological methods, demonstrated that Malaise traps can be successful at recovering lepidopteran biodiversity (Bengtsson, 2024). Our findings corroborate the latter notion. The IBA survey sites and trap locations were not selected to optimize the quantity or diversity of the Lepidoptera catch. Nonetheless, we detected 57% of the natural resident species in a single year. Compared to traditional data, despite the popularity of Lepidoptera, the IBA survey provided better species and occurrence data coverage of sparsely populated regions, and of small and inconspicuous species. Further, the IBA data represents systematically collected sample-based data, which is considerably more powerful in documenting population densities and monitoring long-term trends than traditional data mostly reported ad hoc. Validation of selected new discoveries with full-length individual-specimen barcoding also suggested that the data is reliable. Discoveries include seven putative new species, four species new to the country, and two species occurring well beyond their known ranges. Below, we discuss each of these findings in turn.

The recovery of a large fraction of the lepidopteran fauna attests to two considerations: first, that Malaise traps offer an efficient method for sampling Lepidoptera, and second, that even a single year of high-throughput sampling is sufficient for characterizing a major part of insect biodiversity. Finding more than half of the nation’s lepidopteran diversity (1535/2990 species) in this single sampling effort should be put in a perspective: first, our characterisation of the national fauna was based on a mere 197 traps at 100 sites in a sparse grid (Fig. 1), and second, the point of comparison is to centuries of tireless work by a large lepidopterist community. The 1,535 species (not including new discoveries) detected in the IBA sampling bout compares even more favorably to the 2,379 species recorded by the entire lepidopterist community of Sweden during 2019; it represents two-thirds of it. Importantly, the latter number draws on countless hours invested in all possible sampling methods from visual surveys to light traps, pheromone traps, bait traps, and photography.

The proportion of the national species pool recorded by high-throughput sampling was distributed fairly evenly across lepidopteran families as well as between the traditionally recognised groups of Micro– vs Macrolepidoptera (Fig. 2). In other words, the constancy occurred irrespective of large variation in size, wing load, body plan, life history and diurnal activity. This is a hope-inspiring pattern, since it provides evidence against the notion that Malaise traps would disproportionately sample some taxa at the expense of others. Indeed, it elevates Malaise traps to a key method for sampling Lepidoptera. Admittedly, few lepidopterists will be enthusiastic about the ethanol-soaked specimens generated by this type of sampling (but Schmidt et al. 2019 offer instructions on how to make the most of this type of collection). Nonetheless, Malaise traps yield key metrics of the local lepidopteran fauna – with insights well beyond those provided by a traditional insect collection or a photo collection.

While the data collected by lepidopterists tends to be biased towards large and conspicuous, preferably diurnal species occurring in densely populated areas of the country and popular summer destinations, the high-throughput survey data appears to provide a more even taxonomic and regional coverage (Figs. 3–4). Interestingly, the IBA data indicate that the species richness of local Lepidoptera communities (alpha-diversity) represents a high proportion of the regional species pool (gamma-diversity), resulting in low beta-diversity (Whittaker, 1972). This suggests an interesting ecological pattern, i.e. low species turnover (i.e., low beta-diversity) of lepidopteran communities. This pattern contrasts with that of other taxa, such as plants and fungi (De La Peña Aguilera, 2024), and reveals a key dimension of biodiversity patterns.

Metabarcoding is known to be associated with numerous data quality issues. Deviant sequences, which appear to represent distinct species, may instead reflect variation at the population level, heteroplasmy – presence of several mitochondrial haplotypes within one individual (Kang et al., 2016; Magnacca & Brown, 2010; Ricardo et al., 2020) – or nuclear sequences of mitochondrial origin (NUMTs) (Song et al., 2008). Most NUMTs are short and include indels and/or premature stop codons (IPSCs) (Hebert et al., 2023), which renders them easy to identify and filter out. However, some NUMTs do not have IPSCs and are therefore particularly difficult to distinguish from genuine COI sequences. This highlights the need for caution when unknown COI haplotypes are discovered in metabarcoding studies. In addition, contamination, chimera formation during PCR (Bjørnsgaard Aas et al., 2017), and sequencing errors can contribute to spurious clusters being detected in metabarcoding studies. The extensive congruence between the IBA survey and traditional Lepidoptera data, and the high success rate of the validation tests using full-length barcoding of individual specimens, suggest that our bioinformatic processing successfully tackled those issues and the resulting data is reliable. Undoubtedly, this is partly explained by the completeness of the Lepidoptera reference library, which was used in quality filtering of the data, and the relatively long amplicon sequence targeted (418 bp), increasing the accuracy of library matching. Admittedly, data for insect clades less well-represented in databases (i.e. Diptera, Hymenoptera – Ronquist et al., 2020) or with greater incidence of NUMTs (Orthoptera, Isoptera – Hebert et al., 2023) could be less precise, and likely more challenging to validate. A completely different type of concern is that variation in PCR primer sites may cause detection failure in DNA metabarcoding (Brandon-Mong et al., 2015). However, an independent analysis of 40 IBA samples at the genus level using PCR-free metagenomic methods generated results that exactly matched the Lepidoptera metabarcoding data (López Clinton et al., 2024), suggesting that this is a minor problem, at least in the Lepidoptera.

The validated new discoveries attest to the power of high-throughput surveying in providing novel information about species representing a cross-section of taxonomic and biological diversity (Fig. 5A-D). IBA detected three species new to Sweden. Among those, *Eupithecia groenblomi*, a small, scarce, and hard-to-identify species in the family Geometridae, which was previously known from both Finland and Norway. Its host plant, European goldenrod (*Solidago virgaurea*), is widely distributed in Sweden. Thus, its previous absence from the Swedish checklist will likely reflect an oversight by Swedish lepidopterists. Another species, *Phyllonorycter mespilella,* is an inconspicuous member of the family Gracillariidae. It shares its food plant with several species that are extremely similar to one another and thus can be easily overlooked even by specialists. Molecular data, on the other hand, distinguishes between those species with ease. Finally, *Agriopis budashkini*, is a surprising find in Sweden, since its known range is Southern Europe, Balkans and Crimea. However, it is likely a cryptic species of which the entire range is unknown, since it is extremely easily confused with *Agriopis aurantiaria*. More speculatively, this cryptic species might have spread northwards already over a decade ago, and be the hidden player behind reports of outbreaks of the geometrid moth known as *A. aurantiaria* in northern Norway (Jepsen et al., 2011). An alternative explanation is that this observation results from mitochondrial introgression from *A. budashkini* to *A. aurantiaria*, a phenomenon documented in several other species of European Lepidoptera (Hausmann et al., 2011). Those possibilities should be investigated further by specialists.

In terms of species-specific ranges, the high-throughput samples revealed records of species outside of their previously-presumed ranges. The two validated range expansions represent slightly different cases. *Aleimma loeflingiana* (Tortricidae) is a leafroller moth that mainly feeds on oak (Kimber, 2007). Most of the IBA records match well the known distribution (GBIF records), but one record stands out. IBA recorded *A. loeflingiana* from the Torneträsk area, in far northern Sweden, way outside of its currently known range (Fig. 5E). This species is confined to oak and despite the high latitude, oak does occur in the vicinity of the trap site, it is therefore possible that the species occurs there. However, the record could also represent a stray vagrant. The other validated example, *Coleophora juncicolella* (Coleophoridae), is a small case-bearing moth, which feeds on common heather (*Calluna vulgaris*) and bell heather (*Erica cinerea*). It is not recorded very commonly in GBIF, and mainly from southern Sweden. The IBA data adds many data points beyond the known species distribution, showing that *C. juncicolella* is in fact widespread throughout the country, as are the host plants (Fig. 5F). The species appears very likely to be a resident species overlooked in the northern part of its range.

Perhaps most impressively, we were able to validate the presence in Sweden of eight new COI variants, which were at least 3% distant from known COI sequences in existing reference libraries. Given the high coverage of lepidopteran taxa in these reference libraries (41% of all insect species globally; boldsystems.org) and reliable bioinformatic clean-ups, this suggests that the new variants are reasonably likely to represent species new to science. A case supporting this conclusion is *Acrocercops andreneli*, detected as a putative new species in the IBA survey (2019), and described in 2023 (Nel et al. 2023). Thus, surprisingly, less than 200 sampling locations from a nation the size of 450,000 km^2^ appears to be sufficient to discover a number of new species in one of the best-surveyed insect faunas of the world.

Overall, our findings point to the massive potential for insect biodiversity surveying and discovery through high-throughput methods, and to equally massive knowledge gaps among the few insect groups thought to be well known. To gain reliable knowledge of the fauna, we need to supplement the seminal collection of biological records by nature enthusiasts with systematic sampling designs generating reliable data in time and space. Needless to say, the two approaches should be seen as complementary rather than alternative to each other. Nonetheless, the massive insights generated by a single year of high-throughput sampling suggests that such methods should be added to any national tally of biodiversity. As a key bonus, these methods are based on *standardized sampling* with a *known sampling effort*, thus massively facilitating the downstream analysis of trends and drivers affecting ecosystems and the services they provide.

## Acknowledgements

This work was supported by the Knut and Alice Wallenberg Foundation (grant KAW 2017.088 to FR), the Swedish Research Council (grant 2018-04620 to FR, grant 2021-03784 to AJMT, grant 2021-05563 to AFA), the Polish National Agency for Academic Exchange (grant PPN/PPO/2018/1/00015 to PL), the Polish National Science Centre (grant 2018/31/B/NZ8/01158 to PL), the European Research Council (Synergy Grant 856506 to TR), the Swedish University of Agricultural Sciences (Career Support grant to TR), the Science for Life Laboratory (SciLifeLab) & Wallenberg Data Driven Life Science Program funded by Knut and Alice Wallenberg Foundation (grants: KAW 2020.0239 and KAW 2017.0003), the National Bioinformatics Infrastructure Sweden (NBIS) at SciLifeLab. The computations and data handling were enabled by resources in projects SNIC 2020/15-307, SNIC 2022/5-211, NAISS 2023/5-209, and NAISS 2024/5-207 (compute) and SNIC 2020/16-248 (storage), provided by the National Academic Infrastructure for Supercomputing in Sweden (NAISS) and the Swedish National Infrastructure for Computing (SNIC) at Uppsala Multidisciplinary Center for Advanced Computational Science (UPPMAX), partially funded by the Swedish Research Council (grant 2022-06725 and 2018-05973). The authors also acknowledge support from the National Genomics Infrastructure at SciLifeLab Solna.

## Data and code availability

Figshare repository with GBIF data – https://figshare.com/account/home#/projects/224766

Github repository with data, code and scripts for the annotation analysis, the analysis of the GBIF data, and for generating the figures in the paper: https://githttps.com/insect-biome-atlas/paper-lep-efficiency

