## Supplementary for "High-throughput biodiversity surveying sheds new light on the brightest of insect taxa"

**Electronic Supplementary Material for**

**Main article DOI:**TBD

**Table of contents:**


**Supplementary Methods S1** **2**
- Analysis of DynTaxa misses 2
- Analysis of BOLD annotation problems 14
- Analysis of unidentified clusters 20
- Analysis of unclassified clusters 33
- Analysis of range expansions 45

**Supplementary Results S1** **45**
- Species potentially new to Sweden 45
- Species potentially new to science 46
- Potential range expansions within Sweden 50

**Supplementary figures** **51**

**Tables** **55**

**References** **55**

### Supplementary Methods S1

We analyzed and complemented the taxonomic identification of all clusters by running a set of manual analyses focusing on clusters that could not be matched to Swedish species in the official DynTaxa checklist. We also analyzed OTU occurrence records that represented significant expansions of the known distribution range of Swedish species. These analyses, detailed below, were followed by selection of some clusters (highlighted in yellow below) for validation using full-length barcoding of individual specimens pulled from the corresponding samples. All queries, responses and R scripts used in the analyses of cluster annotations, as well as the output files, are available in the github repository associated with the paper. The updated taxonomic annotations of OTUS are summarized in the output file “summary_manual_otu_annotations.tsv”.

#### Analysis of DynTaxa misses

We analyzed all species annotations of ASVs in our data, which did not match a species recorded in DynTaxa as reproducing in Sweden (“Bofast och reproducerande” in the DynTaxa database). Specifically, we matched the species-level annotations from our pipeline with the DynTaxa database using the online match functionality at https://www.dyntaxa.se/Match/Settings/214225 on 2023-06-09 at 10:24 CEST, and recorded the Swedish occurrence status if there was a match to DynTaxa. There were 68 species annotations, which did not match a species recorded as reproducing in Sweden in DynTaxa. To assist in the interpretation of these cases, the corresponding sequences were matched to BOLD through the bold package in R, and to GenBank using the online nucleotide BLAST functionality at https://blast.ncbi.nlm.nih.gov/Blast.cgi. Occurrence records were also downloaded from GBIF (https://gbif.org). These searches were completed 2023-06-09–10. The results are summarized below.

*Olethreutes glaciana* [**Tortricidae_cluster9**]

The BIN includes 324 records of this species but also 25 records of *Phiaris bipunctana*, which may be synonymous with *Olethreutes glaciana*. North American lepidopterists use an older generic classification, which results in a different generic name being used in BOLD for these species. Finnish *P. bipunctana* share the same BIN as *O. glaciana*. This OTU is almost certainly *P. bipunctana*, which is recorded from Sweden.

*Thyraylia nana* [**Tortricidae_cluster49**]

This species is apparently referred to in DynTaxa as *Cochylis nana*, which is recorded from Sweden.

*Acleris hippophaeana* [in **Tortricidae_cluster148**]

This species is represented by one ASV that is included in a swarm cluster that also includes four ASVs annotated as *Acleris hastiana*. The single ASV annotated as *A. hippophaeana* has 11,785 reads in 11 samples, while the four *A. hastiana* ASVs have a total of 15,485 reads in 18 samples. There are only four samples in which the two co-occur, and the read ratio in those samples varies highly. The *A. hippophaeana* BOLD BIN is unique to this species in BOLD, but there is some variation in the BINs of *A. hastiana*, and the difference to the nearest neighbor from the *A. hippophaeana* BIN is only 1.92% (p-dist) away; it is the BIN matching European *A. hastiana*. All the *A. hippophaeana* samples are from around Vänern, Vättern and Mälaren in central Sweden while the *A. hastiana* samples are more widely distributed over Sweden. The ASV annotated as *A. hippophaeana* could represent a previously unknown intraspecific barcode variant in *A. hastiana*, or potentially *A. hippophaeana* could be overlooked in Sweden.

Detailed analysis of the BOLD data does not provide conclusive evidence. The *A. hippophaeana* BIN (BOLD:ABZ6730) matching our ASV is represented by a single publicly accessible specimen on BOLD. It is collected in Germany and is associated with a note “ID to be verified”. The picture of the voucher looks like a typical *A. hastiana*, so it may be a misidentification, but the two species are extremely variable and hard to tell apart. In Europe, the CO1 barcode sequences of *A. hastiana* and *A. hippophaeana* are tangled (Mutanen et al. 2016), so the deviating ASV in the IBA data may well represent a previously unrecorded variant of *A. hastiana*. Another possibility is that there is a third species hidden in this complex.

*Olethreutes heinrichana* [**Tortricidae_cluster192**]

This is likely *Phiaris heinrichana*, which is recorded from Sweden.

*Cochylichroa atricapitana* [**Tortricidae_cluster266**]

Apparently, this is synonymous with *Cochylis atricapitana* (Stephens, 1852), which is recorded from Sweden.

*Crambus nemorella* [**Crambidae_cluster2**]

The BIN annotation in BOLD has changed to *Crambus lathoniellus*, which is recorded from Sweden.

*Agriphila tristellus* [**Crambidae_cluster6**]

The BIN annotation in BOLD has been corrected to *Agriphila tristella*, which is recorded from Sweden.

*Catoptria margaritellus* [**Crambidae_cluster9**]

The BIN annotation in BOLD has been corrected to *Catoptria margaritella*, which is recorded from Sweden.

*Scoparia cembrella* [**Crambidae_cluster30**]

The BIN annotation in BOLD has changed to *Scoparia subfusca*, which is recorded from Sweden.

*Platytes cerusella* [**Crambidae_cluster40**]

The BIN annotation in BOLD has been corrected to *Platytes cerussella*, which is recorded from Sweden.

*Eupithecia niphadophilata* [in **Geometridae_cluster9**]

Clustered by swarm with *E. pusillata*, which is the resolved species annotation for this OTU. Seems likely to be a rare CO1 variant of *E. pusillata*, not previously recorded in Europe.

*Euphyia intermediata* [**Geometridae_cluster140**]

This is a North American species but the BIN also has European members annotated as *Euphyia unangulata*. This swarm cluster likely represents the latter, which occurs in Sweden.

*Thera cembrae* [**Geometridae_cluster155**]

This appears to be *Thera obeliscata*. This species has two BINs in BOLD, the most commonly sequenced one is this BIN. The same BIN has also been annotated less commonly as *Thera cembrae*, and *Thera variata*. GBIF should not have resolved the BIN name to *Thera cembrae* according to current data, only to *Thera* sp.

*Eupithecia groenblomi* [**Geometridae_cluster181**]

This OTU is represented by 216 reads in 1 sample. The BIN is unique for this species (4 BOLD records, all from Finland). The species is currently recorded from Finland and Norway but not Sweden. This is a species that is likely to occur in Sweden, and it has been a mystery why it has not been observed yet. Possibly it is because it is rather rare and not easy to find.

*Ectropis excellens* [**Geometridae_cluster183**]

The BIN is unique for this species, which is Asian. The nearest BIN in BOLD is another BIN annotated to the same species. *E. crepuscularia* has two very distinct barcodes in Europe, and these two are also retrieved in the IBA data (Geometridae_cluster47, Geometridae_cluster50). The two barcodes have been studied with genomics means (MM, unpublished data) and they do seem to belong to the same species. However, this OTU seems to be something different, as it is placed in a separate swarm cluster. *E. excellens* is very similar to *E. crepuscularia*, so it could possibly have been overlooked. However, the OTU occurs in only 4 samples, always together with *E. crepuscularia*. The read ratios (45.2,1084.1) suggest that this OTU could be a numt of *E. crepuscularia*, assuming that the latter has two distinct CO1 variants (2 of the samples with assumed numts are dominated by one CO1 variant, the other 2 samples by the other CO1 variant of *E. crepuscularia*).

*Agriopis budashkini* [**Geometridae_cluster186**]

The BIN is unique for this species (three BOLD records, two from identified individuals and one from an unidentified individual from the same general area - Greece). The females of this species are wingless. The species is recorded in GBIF from Yugoslavia, Greece, Cyprus and Crimea.

There is also a record from Hungary, which has been disputed (László & Volynkin 2023). It would be surprising if the species occurred in Sweden, but it could be overlooked because of the similarity to *A. aurantiaria*. Interestingly, the latter species is placed in a separate swarm cluster in the IBA data (Geometridae_cluster23). There is no co-occurrence with the current OTU, which occurs in one sample with 90 reads. *Agriopis budashkini* was described only in 2009, and it seems plausible that its entire distribution is not known. It is also possible that the two species share barcodes, perhaps due to introgression.

*Xestia speciosa* [**Noctuidae_cluster35**]

This is likely the species that has been referred to as *Xestia arctica* in Sweden.

*Xestia smithii* [**Noctuidae_cluster43**]

This is likely *Xestia baja*, which occurs in Sweden, and shares BIN with *X. smithii*.

*Hyppa contrasta* [in **Noctuidae_cluster55**]

The swarm cluster only includes one ASV annotated to species and BIN, the one annotated to *H. contrasta*. The other ASVs are only annotated to genus (*Hyppa*). The specific BIN identified in the species-level annotation is completely dominated in BOLD by *Hyppa contrasta*, a North American species, but it does also include one member of *Hyppa rectilinea*, which is recorded from Sweden. Most of the *H. rectilinea* specimens in BOLD are in a neighboring BIN, but strangely enough, Sintax does not match most of the Swedish ASVs unequivocally to that BIN. Nevertheless, this OTU seems likely to represent *Hyppa rectilinea*.

*Lacanobia atlantica* [in **Noctuidae_cluster60**]

This is a North American species, which does not share BIN with any other species (23 BOLD records from Canada, 3 from China). In the IBA data, however, there are only 18 reads from a single sample annotated as *L. atlantica*, and they are grouped in the same swarm cluster as ASVs identified as *L. suasa* and *L. thalassina*, both of which are recorded from Sweden. There are two more *Lacanobia* OTUs in the IBA data, and they clearly seem to be *L. oleracea* and *L. w-latinum*. The OTU with mixed annotations is not resolved according to the IBA pipeline. It could represent *L. suasa*, *L. thalassina*, or a mix of the two. The ASV annotated as *L. atlantica* seems likely to be a rare variant of one of these two species, a variant that has not been recorded previously.

*Xestia rhomboidea* [**Noctuidae_cluster114**]

The BOLD BIN annotation has changed to *X. stigmatica*, which is recorded from Sweden.

*Xestia homogena* [**Noctuidae_cluster154**]

This is a Nearctic species, which is closely related to and shares BIN with *Xestia rhaetica*, which is recorded as Swedish. This OTU is likely to be *Xestia rhaetica*, which is also called *Xestia fennica* or *Xestia rhaetica fennica*.

*Oxypteryx atrella* [**Gelechiidae_cluster32**]

In DynTaxa, this species is referred to as *Eulamprotes atrella*, which is recorded from Sweden.

*Oxypteryx unicolorella* [**Gelechiidae_cluster38**]

In DynTaxa, this species is referred to as *Eulamprotes unicolorella*, which is recorded from Sweden.

*Apodia martinii* [**Gelechiidae_cluster53**]

This OTU matches a unique BIN for *A. martinii* (specimens from Åland and central Europe), which is quite distant (6.16% p-dist) from another BIN, annotated as *A. bifractella* (from central and southern Europe). The two species were split recently, and DynTaxa probably incorrectly refers the Swedish records to *A. bifractella* although they actually belong to *A. martinii*.

*Aproaerema taeniolella*, *A. cinctella*, *A. karvoneni* [**Gelechiidae_cluster54**, **Gelechiidae_cluster61**, **Gelechiidae_cluster92**]

In DynTaxa, these species are placed in *Syncopacma*, explaining the match failure. They are all recorded from Sweden.

*Oxypteryx wilkella* [**Gelechiidae_cluster112**]

In DynTaxa, this species is referred to as *Eulamprotes wilkella*, which is recorded from Sweden.

*Oxypteryx superbella* [**Gelechiidae_cluster113**]

In DynTaxa, this species is referred to as *Eulamprotes superbella*, which is recorded from Sweden.

*Monochroa palustrellus* [**Gelechiidae_cluster124**]

In DynTaxa, this species is referred to as *Monochroa palustrella*, which is recorded from Sweden.

*Vanessa atalanta*, *V. cardui* [**Nymphalidae_cluster6**, **Nymphalidae_cluster7**]

Common migrants, temporarily reproducing in Sweden.

*Agonopterix rubidella* [**Elachistidae_cluster33**]

In DynTaxa, this species is referred to as *Agonopterix angelicella*, which is recorded from Sweden.

*Acrolepia pygmaeana* [**Glyphipterigidae_cluster2**]

The BOLD annotation of this BIN has changed to *Acrolepia autumnella*, which occurs in Sweden.

*Adela viridella* [**Adelidae_cluster1**]

The BOLD annotation of this BIN has changed to *Adela reaumurella*, which occurs in Sweden.

*Nemophora scopolii* [in **Adelidae_cluster7**]

Only 60 reads in 2 samples representing one ASV match a BIN that is annotated to this species, which was recently split from the closely related *N. degeerella*. The swarm clustering groups the

*N. scopolii* reads with many ASV reads annotated as *Nemophora degeerella* and a few unclassified *Nemophora* reads. This could either represent unrecorded BIN sharing, or potentially document that both species occur in Sweden, in which case *N. scopolii* is a new record. Another possibility is that the ASV that is annotated as *N. scopolii* is actually a numt of *N. degeerella*, as the latter species is also present and more abundant in the two samples containing the former.

*Nemophora bellella* [**Adelidae_cluster14**]

The BOLD annotation of this BIN has been corrected to *Nemophora bellela*, which is recorded from Sweden.

*Yponomeuta vigintipunctata* [**Yponomeutidae_cluster17**]

The BOLD annotation of this BIN has changed to *Yponomeuta sedella*, which occurs in Sweden (given as *Y. sedellus* in DynTaxa)

*Euhyponomeutoides rufella* [**Yponomeutidae_cluster19**]

The BOLD annotation of this BIN has changed to *Euhyponomeutoides albithoracellus*. This species is recorded from Sweden.

*Phyllonorycter lautella* [**Gracillariidae_cluster38**]

In DynTaxa, the species name is spelled *Phyllonorycter lautellus*. The species is recorded from Sweden.

*Phyllocnistis populiella* [**Gracillariidae_cluster58**]

This is a North American species that shares BIN with *P. labyrinthella*. This OTU in the IBA data presumably represents the latter, which occurs in Sweden.

*Phyllonorycter mespilella* [**Gracillariidae_cluster61**]

There are 1,227 reads in one sample of this OTU. In DynTaxa, the species name is spelled *Phyllonorycter mespilellus*. The species has not been recorded from Sweden but it occurs in

Denmark. The BIN is unique to this species. There are 1,265 IBA samples with *Phyllonorycter* OTUs in them in the IBA data. However, the sample containing the current OTU has no other *Phyllonorycter* OTUs in it. This is a species for which overlooking is easy. It is extremely similar to several other species that share the same food plants. This could potentially represent a new species for Sweden; the species occurs close to where the OTU was found, and the food plant is present there. Barcode variability of this group has been studied intensively (Lopez-Vaamonde 2021), suggesting that barcode sharing is unlikely.

*Phyllonorycter klemannella* [**Gracillariidae_cluster67**]

In DynTaxa, the species name is spelled *Phyllonorycter klemannellus*. The species occurs in Sweden.

*Phyllonorycter cerris* [in **Gracillariidae_cluster84**]

This swarm cluster contains two ASVs, one unclassified *Phyllonorycter* and one matching a BIN that is unique to *Phyllonorycter cerris*. The latter BIN is closest to *P. quercusmacrolepis*. Both of these species occur in central to southern Europe. This OTU seems likely to represent something else. The two ASVs BLAST to *P. quercifoliella* (98.33% and 98.56% identity, respectively). They are represented by few reads (7 reads in 1 sample, and 6 reads in 1 sample). In the same samples, *P. quercifoliella* is present with 2,736 and 1,734 reads, respectively. The latter values are in the 85-90% quantile of reads for this species, that is, this OTU pops up in samples where we have a lot of reads of *P. quercifoliella*. Therefore, this OTU seems likely to represent a numt in *P. quercifoliella*.

*Dendrolimus superans* [**Lasiocampidae_cluster4**]

The BIN matching this cluster is also the only BIN associated with *Dendrolimus pini*. This OTU in the IBA data is likely to represent the latter, which is a common Swedish species.

*Poecilocampa alpina* [**Lasiocampidae_cluster5**]

Most of the specimens belonging to this BIN are annotated in BOLD as *Poecilocampa populi*. This OTU is likely to represent the latter, which is a common Swedish species.

*Trichiura crataegi* [**Lasiocampidae_cluster7**]

This is a simple matching problem in DynTaxa, apparently due to an erroneous entry in the latter. This species occurs in Sweden.

*Taleporia politella* [**Psychidae_cluster3**]

The BOLD annotation of this BIN has changed to *T. tubulosa*, which is a Swedish species. There are two BINs for *T. tubulosa*. Both BINs are represented in the IBA data, and they are grouped in the same swarm cluster.

*Spindasis takanonis* [**Lycaenidae_cluster13**]

The BOLD annotation of this BIN has changed to *Satyrium w-album*, which is a Swedish species.

*Gillmeria ochrodactyla* [**Pterophoridae_cluster9**]

In DynTaxa, this species is referred to as *G. tetradactyla*, which is a Swedish species.

*Mompha decorella* [**Momphidae_cluster5**]

The BOLD annotation of this BIN has changed to *M. divisella*, which is a Swedish species.

*Fomoria weaveri* [**Nepticulidae_cluster1**]

In DynTaxa, this species is referred to as *Ectoedemia weaveri*. This species occurs in Sweden.

*Stigmella nylandriella* [**Nepticulidae_cluster5**]

This is a simple matching problem in DynTaxa, apparently due to an erroneous entry in the latter. This is a Swedish species.

*Zimmermannia atrifrontella* [**Nepticulidae_cluster26**]

In DynTaxa, this species is referred to as *Ectoedemia atrifrontella*. This species occurs in Sweden.

*Fomoria septembrella* [**Nepticulidae_cluster46**]

In DynTaxa, this species is referred to as *Ectoedemia septembrella*. This species occurs in Sweden.

*Zimmermannia amani* [**Nepticulidae_cluster57**]

In DynTaxa, this species is referred to as *Ectoedemia amani*. This species occurs in Sweden.

*Etainia albibimaculella* [**Nepticulidae_cluster71**]

In DynTaxa, this species is referred to as *Ectoedemia albibimaculella*. This species occurs in Sweden.

*Etainia sericopeza* [**Nepticulidae_cluster76**]

In DynTaxa, this species is referred to as *Ectoedemia sericopeza*. This species occurs in Sweden.

*Drepanidae_Achlya flavicornis* [**Drepanidae_cluster4**]

This is a simple error in the GBIF resolution of the species annotation. This OTU is *Achlya flavicornis*, which is a Swedish species.

*Pseudoips prasinana* [**Nolidae_cluster8**]

In DynTaxa, the species name is spelled *Pseudoips prasinanus*. This species occurs in Sweden.

*Batrachedra confusella* [**Batrachedridae_cluster3**]

A recently described species (Berggren et al. 2022), not recorded from Sweden yet but likely to occur in the country. The two species of the genus that are currently recorded from Sweden are also represented in the IBA material: *B. praeangusta* (Batrachedridae_cluster1) and *B. pinicolella* (Batrachedridae_cluster2). All three species/clusters are represented in multiple samples and with a decent number of reads, with no co-occurrence, as follows: 103,204 reads in 57 samples (cluster 1), 21,066 reads in 12 samples (cluster 2), and 7,027 reads in 8 samples (cluster 3). Thus, this seems a likely new species record for Sweden.

*Dioryctria mutatella* [**Pyralidae_cluster9**]

The BOLD annotation of this BIN has changed to *Dioryctria simplicella*, which is a Swedish species.

*Plodia interpunctella* [**Pyralidae_cluster21**]

This is a synanthropic species, regularly recorded from Sweden.

*Dyseriocrania subpurpurella* [**Eriocraniidae_cluster2**]

This is a simple matching problem in DynTaxa, apparently due to an erroneous entry in the latter. This is a Swedish species.

*Epermenia chaerophyllellus* [**Epermeniidae_cluster1**]

The BOLD annotation has been corrected to *Epermenia chaerophyllellla*, which is also the name form used in DynTaxa. This is a Swedish species.

*Prays curtisella* [**Praydidae_cluster1**]

The ASV in the IBA data annotated by Sintax as *Prays curtisella* belongs to a swarm cluster of two ASVs matching two different BINs. The BIN formerly annotated as *P. curtisella* is now annotated as *P. ruficeps* in BOLD. The annotation of the other BIN is either *P. oleae* or *P. fraxinellus* (mix of specimens). Both *P. ruficeps* and *P. fraxinellus* are Swedish species, and it is possible that they are mixed in this OTU. The two ASVs BLAST to *P. ruficeps* and *P. fraxinellus*, respectively (with close matches to other *Prays* species in this triplet).

*Bedellia somnulentella* [**Bedelliidae_cluster1**]

Recorded from Sweden but unknown whether it is reproducing.

#### Analysis of BOLD annotation problems

We then analyzed clusters that were resolved by our pipeline to a unique BIN, but not to a unique species annotation. This could be due to the GBIF name resolution algorithm being overly cautious, as it did not resolve the name of BOLD BINs unless 80% or more of the records were identified to species and had exactly the same species name. It could also be due to a range of other causes. In total, there were 185 Lepidoptera OTUs of this kind in the IBA data. Before more thorough analysis, we updated the species annotation of these OTUs to the most frequent species-level annotation in the most up-to-date version of BOLD (using the "bold" package in R, performed on 2023-06-11), and then matched the updated annotation to DynTaxa to find if the species were recorded from Sweden. We focus the detailed analysis below on the 27 OTUs that could not be matched to Swedish species using this procedure (the first 27 clusters). We also looked at the five OTUs that were annotated using this procedure to species, which had been previously associated with other OTUs in the IBA data (the last five clusters). Note that we did not look at 17 OTUs, which had "Unspecified*" as their updated BIN annotation.

The detailed analysis was based on online searches in BOLD, GenBank and DynTaxa performed on 2023-06-11—12. DynTaxa matches were performed using the online match functionality on 2023-06-11 at 15:47 CEST. Detailed data underlying this analysis are provided in the file "bold_annotation_problems.tsv". The name of each cluster represents the closest match in GenBank.

**Various causes**. The first six clusters are associated with different types of annotation problems, as follows.

**Nepticulidae_cluster45**: *Stigmella salicis* cluster 2

**Nepticulidae_cluster59**: *Stigmella salicis* cluster 6

**Nepticulidae_cluster65**: *Stigmella salicis* cluster 3

Apparently, there is some variation within this species. In BOLD, cluster 2 is fairly widespread and has been recorded from Finland and Norway, among other countries. Cluster 6 is only recorded from the UK, and cluster 3 from Finland, Norway and France. In the Swedish IBA data, we have two ASVs belonging to cluster 2, and one each belonging to cluster 6 and cluster 3. These could potentially represent intra-specific variation or cryptic species.

**Ypsolophidae_cluster14**: *Ochsenheimeria* sp.

Dyntaxa lists four species in the genus *Ochsenheimeria* as Swedish: *O. mediopectinella*, *O. taurella*, *O. urella* and *O. vacculella*. Interestingly, there are four OTUs in the genus in the IBA data (Ypsolophidae_cluster 9, 12, 14 and 16). These clusters correspond perfectly to four different BINs. Three of these BINs have been encountered in Finland, the remaining one only in Austria. Two of the BINs are annotated by Sintax (through GBIF) as *O. mediopectinellus* but the BOLD annotation of these two BINs has since changed to *O. urella*. One BIN is annotated as *O. vacculella* and the remaining one (the one only recorded from Austria) as *Ochsenheimeria* sp. (the one corresponding to Ypsolophidae_cluster14). The latter cluster does not co-occur with any of the other three clusters. The correct identification of the four *Ochsenheimeria* clusters in the IBA data remains uncertain.

**Tortricidae_cluster84**: *Neocochylis dubitana*

This is *Cochylis dubitana*, a species recorded from Sweden.

**Geometridae_cluster73**: *Idaea deversaria*

This appears to be a simple matching problem in DynTaxa due to an older synonym. The species is recorded from Sweden.

**Difficulties separating North American and Palaearctic species**. For the OTUs below, the most frequent annotation in BOLD is shared by less than 80% of the BOLD specimens. It is commonly the case that the most frequent BOLD annotation is for a North American species (or name form), even though the BIN is shared with a closely related European species (or name form).

**Geometridae_cluster51**: *Scopula frigidaria*

The species in the new BOLD BIN annotation has apparently not been recorded from Sweden previously. However, the BIN is shared with *Scopula ternata* and *S. siccata*. This may well be *S.*

*ternata*, which has been recorded from Sweden. The latter species is only represented in this BIN in BOLD. It is also possible that this cluster may correspond to *S. frigidaria*, which supposedly occurs in Fennoscandia even though it has not been recorded from Sweden yet, or a mix of the two species. The OTU is abundant; it occurs in 133 samples with a total of more than 100,000 reads.

**Geometridae_cluster92**: *Eupithecia albicapitata*

This species is North American. The BIN is shared with *E. analoga*, which occurs in Sweden. The latter is only represented in this BIN in BOLD.

**Drepanidae_cluster5**: *Drepana bilineata*

This species is North American but the BIN is shared with *Falcaria lacertinaria*, which occurs in Sweden. The latter is only reported from this BIN in BOLD.

**Sphingidae_cluster4**: *Hyloicus pinastri*

This species is given as *Sphinx pinastri* in DynTaxa, a common Swedish species.

**Noctuidae_cluster165**: *Leucania dia*

This species is North American but the BIN is shared with *Leucania comma*, which occurs in Sweden. The latter is only reported from this BIN in BOLD.

**Noctuidae_cluster192**: *Melanchra pulverulenta*

This species is North American but the BIN is shared with *Ceramica pisi*, which occurs in Sweden. The latter is only reported from this BIN in BOLD.

**Gracillariidae_cluster68**: *Caloptilia semifascia*

The BIN is shared with *C. jurateae*. According to DynTaxa, only the latter occurs in Sweden. This IBA OTU is presumably *C. jurateae*, which is only represented in this BIN in BOLD.

**Noctuidae_cluster98**: *Enargia decolor*

This is a North American species. The BIN is shared with *E. paleacea*, which is recorded from Sweden. The latter is only reported from this BIN in BOLD.

**Hesperiidae_cluster4**: *Pyrgus malvoides*

This BIN is shared between *Pyrgus malvoides* and *Pyrgus malvae*. The latter is a Swedish species.

**Tortricidae_cluster131**: *Pammene perstructana*

This is one of two BINs for a North American species. This BIN is shared with *Pammene clanculana*, which occurs in Sweden.

**Erebidae_cluster47**: *Spilosoma lutea*

This is a matching problem in DynTaxa, where the species is given as *Spilarctia lutea*. This species occurs in Sweden.

**Tortricidae_cluster135**: *Pammene perstructana*

This is the second of two BINs for a North American species (see Tortricidae_cluster131). This BIN is shared with *Pammene obscurana* which occurs in Sweden.

**Tineidae_cluster7**: *Nemapogon koenigi*

This BIN is shared with *Nemapogon wolffiella*, which is a Swedish species (given as *N. wolffiellus* in DynTaxa). Neither species is represented outside this BIN in BOLD.

**Noctuidae_cluster65**: *Papestra quadrata*

The BIN is shared between this species and *P. biren*, which occurs in Sweden. Neither species is represented outside this BIN in BOLD.

**Tortricidae_cluster244**: *Acleris* sp. 4

This BIN is most commonly annotated as *Acleris ferrumixtana* (from Finnish specimens), although there are 7 undetermined *Acleris* specimens placed in the same BIN from Canada. *Acleris ferrumixtana* is only represented in this BIN in BOLD. There is one specimen in this BIN from Norway labeled *A. implexana*, but all other specimens of *A. implexana* in BOLD are North American and are grouped in a different BIN. This BIN is not found in the IBA data. DynTaxa records *A. implexana* as Swedish but does not include *A. ferrumixtana*. There seems to be some name confusion here. It seems that this cluster represents the same species as that named *A. implexana* in DynTaxa and in Norway but as *A. ferrumixtana* in Finland.

**Tortricidae_cluster83**: *Argyrotaenia velutinana*

According to DynTaxa, there is only one species in the genus *Argyrotaenia*, *A. ljungiana*. This species shares this BIN in BOLD with *A. velutinana* and several other species of *Argyrotaenia*. Unlike some of the other species, *A. ljungiana* is only represented in this BIN in BOLD. In total, there are 26 specimens in BOLD of this species from all over Europe, all placed in this BIN.

**Momphidae_cluster13**: *Mompha* sp.

There are 15 Swedish species in this genus in DynTaxa, and 13 OTUs in the IBA data that are annotated to the genus. This OTU is the only one that is not annotated to a Swedish species. It appears to be *Mompha sexstrigella*. There are two specimens in BOLD of *M. sexstrigella* from Europe (Finland), both are placed in this BIN. However, the North American specimens of *M. sexstrigella* are placed in different BINS in BOLD (25 in one BIN, and 4 in another).

**Tortricidae_cluster180**: *Acleris* cf. *emargana*

The annotations of the specimens in BOLD placed in this and related *Acleris* BINs is quite confusing. Some guesswork and BLAST against GenBank suggests that this OTU could be *A. effractana*, which is a Swedish species.

**Elachistidae_cluster24**: *Elachista serricornis*

This appears to be a matching problem in DynTaxa. The species occurs in Sweden.

**Tortricidae_cluster14**: *Olethreutes metallicana*

This species is the same as *Phiaris metallicana*, which occurs in Sweden.

**Pyralidae_cluster5**: *Ortholepis myricella*

This is a North American species. The BIN is shared with *O. rhodorella* and *O. vacciniella*. All three species are restricted to this BIN in BOLD. Only *O. vacciniella* appears to occur outside of North America, and this species is recorded from Sweden.

**Updated annotation matches previous species annotation**. The following five clusters represent those cases where the updated annotation matched a previous species annotation. These might represent heteroplasmy, numts or sequencing errors.

**Eriocraniidae_cluster9**: *Eriocrania semipurpurella*

**Eriocraniidae_cluster3**: *Eriocrania semipurpurella*

European specimens of this species are split into three BINs in BOLD. The same split is partly reflected in the OTU of the IBA data, as follows: Eriocraniidae_cluster3 matches BOLD:AAB3768, Eriocraniidae_cluster4 matches BOLD:AAB3767 and BOLD:AAB3764, and Eriocraniidae_cluster9 matches BOLD:AAB3764. According to BOLD, AAB3764 is 3.23% different from its nearest neighbor, AAB3767, while the latter is closest to AAX8459, annotated as "*Eriocrania* sp. 3". AAB3768 is closest to AAB3767 at 6.25% p-dist. The clusters occur in 3, 17 and 6 samples, respectively. Cluster 3 and 4 occur in one sample together, and then in equivalent numbers (996 and 1,023 reads, respectively). It seems like these three OTUs could potentially represent at least two, possibly three different species.

**Tortricidae_cluster175**: *Epiblema scutulana*

This species is split into two BINS in BOLD (2.56% p-dist). The same split occurs in the IBA data. This OTU matches BOLD:AAC0715, while Tortricidae_cluster286 matches BOLD:AAP7460. Cluster 175 occurs in 2 samples. Cluster 286 only occurs in one of those samples, and with a few reads (13 reads, compared to 4,043 for cluster 175 in the same sample). Potentially, cluster 286 could represent a heteroplasmic variant or numt. Also in BOLD, there are more specimens of BOLD:AAC0715 than of BOLD:AAP7460.

**Geometridae_cluster18**: *Epirrita autumnata*

This would be the third IBA OTU annotated as *Epirrita autumnata*. The three OTUs cover all five BOLD BINs that match European specimens of *E. autumnata* as follows: Geometridae_cluster2 matches BOLD:ACE7803, BOLD:AAA5906 and BOLD:ABY8748; Geometridae_cluster10 matches BOLD:AAA5909; and Geometridae_cluster18 matches BOLD:AAA5907. All three OTUs are common (166, 70 and 82 samples respectively). There is no obvious pattern of co-occurrence or lack of co-occurrence. Further analysis of the distribution and genetics of the three IBA OTUs may help shed some light on the nature of this genetic variation.

**Noctuidae_cluster97**: *Diarsia mendica*

This is the second IBA OTU annotated as *D. mendica*, the other one being Noctuidae_cluster49. They correspond to two different BINs, BOLD:ABZ6600 (cluster 49) and BOLD:AAB0038 (cluster97). Cluster 49 occurs in 22 samples, and cluster 97 in 4 samples. They only co-occur in one sample, and then in equal numbers (2,266 and 3,572 reads, respectively). Unclear whether this represents intra- or interspecific variation; neither OTU appears to be a numt.

#### Analysis of unidentified clusters

The analysis below focuses on the cleaned nochimera swarm clusters (OTUs), which are not resolved to a unique BIN nor to a unique species annotation. In total, there are 83 OTUs of this kind in the cleaned IBA data.

Before analyzing the data, we matched the representative ASV sequence of each cluster to BLAST and recorded the key parameters of the top hit. If the top hit was to a species represented in other IBA OTUs, we looked at the frequency of the OTUs and their overlap to see if the unidentified OTU might be a likely numt. In those cases, we might expect the numt to always co-occur with larger numbers of true CO1 sequences from the species to which it matches most closely. If the percent identity of the BLAST hit was very high, and the hit was to an entry described as a mitochondrial sequence, and the taxon was not previously present in the IBA data, the OTU might represent a species missed by the Sintax annotation for some reason. Finally, if the BLAST hit was to a sequence not marked as mitochondrial in BOLD, the OTU would seem to represent a nuclear copy or a numt of CO1. Below, we briefly summarize the conclusions for all 83 clusters along these lines. The underlying data are in the file “unidentified.tsv”. The file can be regenerated using the scripts “generate_ufo_fasta.R” and “generate_ufo_table.R”, with an intervening submission of the fasta file generated by the first script to the web-based BLAST search interface, and saving the 83 hit table descriptions as separate csv files. The BLAST search used here was performed on 2023-06-12 using default settings at https://blast.ncbi.nlm.nih.gov/.

**Elachistidae_cluster1:** *Elachista exactella*

This appears to be a Sintax annotation failure. Potentially, this is due to an erroneous annotation in BOLD, which has been corrected now.

**Crambidae_cluster3:** *Agriphila straminella*

This appears to be a Sintax annotation failure.

**Gracillariidae_cluster1:** *Acrocercops tacita*

This may well represent an undescribed or unsequenced species. The representative ASV has only 95.44% identity to *A. tacita*, which is a southern European species on oaks. It is about as distant to *A. brongniardella* according to the BOLD tree. There is some variation in *A. brongniardella* ASVs in IBA, causing swarm to split the ASVs into five clusters: Gracillariidae_cluster2, 87, 90, 93 and 98. However, all ASVs in these clusters match the same BOLD BIN, BOLD:AAE7282, which is annotated as *A. brongniardella* in BOLD. There are 107 samples with 359,137 reads in total of *A. brongniardella* clusters and 110 samples with 454,609 reads in total of the current OTU. The two groups only co-occur in 49 samples, and in those samples the read ratio range is wide (0.008, 2916.9). The distributional ranges seem to overlap to a large extent, although it is possible to find Swedish regions where only one of the types is encountered in the IBA data. There seems to be no difference in the habitats where they are encountered; both seem to be widespread across habitats.

**Tortricidae_cluster21:** *Apotomis inundana*

This appears to be a Sintax annotation failure.

**Tortricidae_cluster46:** *Phiaris palustrana*

This appears to be a Sintax annotation failure.

**Lycaenidae_cluster2:** *Plebejus argus*

This cluster matches two different BOLD BINs, one of which is fairly consistently annotated as *P. argus*, the other one of which is annotated with many different species in *Plebejus* but most commonly (about half of 748 specimens) with *P. idas*, which is also a Swedish species. This cluster may well consist of two (or more) species of *Plebejus*.

**Nymphalidae_cluster3:** *Lasiommata megera*

This appears to be a Sintax annotation failure.

**Argyresthiidae_cluster5:** *Argyresthia pruniella*

This appears to be a Sintax annotation failure.

**Geometridae_cluster62:** *Dysstroma latefasciata*

This appears to be a Sintax annotation failure.

**Noctuidae_cluster55:** *Hyppa rectilinea*

This appears to be a Sintax annotation failure. See also the analysis of DynTaxa misses for more information on this cluster.

**Elachistidae_cluster17:** *Elachista humilis*

This appears to be a Sintax annotation failure.

**Noctuidae_cluster60:** *Lacanobia thalassina*

This appears to be a Sintax annotation failure. See also the analysis of DynTaxa misses for more information on this cluster.

**Elachistidae_cluster20:** *Agonopterix alstromeriana*

This appears to be a Sintax annotation failure.

**Lypusidae_cluster4:** *Pseudatemelia flavifrontella*

This appears to be a Sintax annotation failure. The species is given in DynTaxa as *Agnoea flavifrontella*.

**Lypusidae_cluster5:** *Lypusa maurella*

This appears to be a Sintax annotation failure.

**Nepticulidae_cluster8:** *Stigmella myrtillella*

This appears to be a Sintax annotation failure.

**Noctuidae_cluster84:** *Lycophotia porphyrea*

This appears to be a Sintax annotation failure.

**Coleophoridae_cluster23:** *Coleophora otidipennella*

This appears to be a Sintax annotation failure; the cluster has many ASVs matched to the BIN that corresponds to *Coleophora otidipennella* in BOLD.

**Gelechiidae_cluster63:** *Bryotropha terrella*

This seems likely to represent a numt of *B. terrella*.

**Tortricidae_cluster148:** *Acleris hippophaeana*

The representative ASV of this clusters appears to be this species, which has not been encountered in Sweden before. In terms of annotations, the cluster is a mix of *A. hippophaeana* and *A. hastiana*. See the previous analysis of Dyntaxa misses for details.

**Adelidae_cluster11:** *Cauchas fibulella*

At 96.41% identity, this appears somewhat unlikely to be *C. fibulella*. The OTU occurs in 14 samples with 18,467 reads. There are no other IBA OTUs in the genus. This could potentially represent a new or unsequenced species close to *C. fibulella*. On the other hand, *C. fibulella* is a Swedish species that is not encountered in the IBA data, so perhaps it could be this species anyway. However, specimens of this species have been barcoded quite widely, so it would be surprising if the Swedish specimens would regularly be this distinct. One possibility is that the used primers amplify a numt but not the actual barcode of *C. fibulella*. It is also possible that Swedish specimens really have a distinct CO1 sequence, such cases are known.

**Noctuidae_cluster95:** *Apamea rubrirena*

This appears to be a Sintax annotation failure.

**Nepticulidae_cluster13:** *Stigmella lappovimella*

This appears to be a Sintax annotation failure.

**Crambidae_cluster33:** *Catoptria permutatellus*

This appears to be a Sintax annotation failure. The species is given as *C. permutatella* in DynTaxa.

**Coleophoridae_cluster31:** *Coleophora alnifoliae*

This appears to be a Sintax annotation failure.

**Tortricidae_cluster179:** *Eucosma aspidiscana*

This appears to be a Sintax annotation failure but it could also be that this cluster represents a mix of several species of *Eucosma*.

**Coleophoridae_cluster38:** *Coleophora serratella*

This appears to be a resolution error; almost all ASVs are matched to a BIN that corresponds to *C. serratella*. There is another IBA cluster that matches this species (Coleophoridae_cluster 65) but it might represent a rare variant or a numt; the current cluster is the dominant one.

**Nepticulidae_cluster20:** *Stigmella luteella*

This appears to be a Sintax annotation failure.

**Noctuidae_cluster127:** *Euxoa nigricans*

This appears to be a Sintax annotation failure.

**Gracillariidae_cluster47:** *Phyllonorycter oxyacanthae*

This appears to be a Sintax annotation failure.

**Nepticulidae_cluster25:** *Stigmella glutinosae*

This appears to be a Sintax annotation failure.

**Coleophoridae_cluster45:** *Coleophora betulella*

This appears to be a Sintax annotation failure. There is another IBA cluster annotated to the same species, cluster 54, but cluster 45 is the most abundant one. BOLD also splits this species into two BINs, but in another way than the IBA data do.

**Nepticulidae_cluster29:** *Stigmella salicis*

This appears to be a Sintax annotation failure.

**Bucculatricidae_cluster9:** *Bucculatrix notella*

This OTU only has a 94.50% identity to *B. notella*, so it is unlikely to be this species, which is not recorded as Swedish. The OTU is represented by 1,870 reads in 2 samples. There are no other Bucculatricidae OTUs in those samples. The ASV goes in the BOLD-generated tree as a sister to *B. cristatella*, which appears reasonable. *Bucculatrix cristatella* is both genetically and morphologically a variable species, and it is represented by two swarm clusters in the IBA data (Bucculatricidae_cluster1 and Bucculatricidae_cluster2). It cannot be excluded that this cluster represents a genetically distinct CO1 variant in *B. cristatella*, although the distance to other barcoded specimens suggests that it represents a distinct but cryptic species.

**Coleophoridae_cluster47:** *Coleophora otidipennella*

The cluster, represented by 1,806 reads in three samples, only contains a single ASV with 96.89% identity to *C. otidipennella*. Of the three samples, one does not contain any other *Coleophora* OTU, and this is also the sample with the highest number of reads of the current cluster, which is not suggestive of this OTU being a numt in *Coleophora*. This OTU could represent a new or unsequenced species or, alternatively, it does represent *C. otidipennella*, which is recorded from Sweden and represented by Coleophoridae_cluster23 in the IBA data (see above).

**Praydidae_cluster1:** *Prays ruficeps*

This appears to be a Sintax annotation failure.

**Tortricidae_cluster210:** *Eucosma aspidiscana*

At only 96.41% identity, this appears relatively unlikely to be a CO1 sequence of *E. aspidiscana*. It could represent a rare variant or possibly a numt, even though the single sample containing it has 1,570 reads. There are no other *Eucosma* OTUs in that sample. The genus *Eucosma* is unusual as it shows a fair amount of barcode sharing across species.

**Tortricidae_cluster214:** *Philedonides lunana*

At only 96.64% identity, this might appear relatively unlikely to be a CO1 sequence of *P. lunana*. The latter is also represented as a different cluster in the IBA data. Thus, this OTU could possibly represent a new or previously unsequenced species. However, the current OTU is only present in 3 samples, and in a significant number of reads (more than 7 reads) in only one sample. So it could be that this actually is a variant belonging to *P. lunana*, and the samples with few reads could potentially represent contamination. The read numbers are about the same in the sample where the two co-occur, suggesting that, if it does indeed belong to the same species, this is a CO1 variant rather than a numt.

**Geometridae_cluster164:** *Yezognophos vittaria*

This appears to be a numt of *Elophos vittaria*, which is represented by Geometridae_cluster13. Cluster 164 occurs in 39 samples, always together with and in lower numbers than cluster 13.

**Tortricidae_cluster218:** *Epinotia crenana*

At only 92.58% identity, this appears unlikely to be a CO1 sequence of *E. crenana*. The latter is represented as a different cluster in the IBA data, and there is no sample co-occurrence. This could possibly represent a new or previously unsequenced species. It is present in 1,569 reads in 2 samples. There is a co-occurring *Epinotia* OTU in one of the two samples, but in the one containing the smallest number of reads of the current OTU.

**Tortricidae_cluster219:** *Orthotaenia undulana*

This appears to be a numt of *O. undulana*.

**Tortricidae_cluster234:** *Archips betulanus*

At 96.65% identity, this could perhaps be *Archips betulana*, which is a Swedish species. The species is not recorded otherwise in the IBA data.

**Elachistidae_cluster59:** *Elachista zernyi*

This appears to be a Sintax annotation failure.

**Tortricidae_cluster243:** *Epinotia signatana*

This sequence matches different *E. signatana* sequences at 100.0% and 94.96% similarity, respectively. The IBA data suggests that this OTU may be a numt of *E. signatana*.

**Noctuidae_cluster168:** *Mythimna impura*

This appears to be a numt of *M. impura*.

**Tortricidae_cluster248:** *Notocelia roborana*

At 95.69% identity, this may be unlikely to represent *N. roborana*. The small number of reads may also suggest that the OTU could represent noise. However, the beginning and the end of the sequence BLAST with similar results, suggesting that it is not a chimera. Also, there is no sample co-occurrence with any other OTU annotated as *Notocelia* in IBA (there are six of them), suggesting that it is not a numt. Interestingly, there are seven species in the genus recorded from Sweden. Five of them are represented in the other IBA data, but the species known as *N. roborana* is missing (as is *N. tetragonana*). *N. cynosbatella* is split into two in the IBA data, in BOLD it is split into three BINs. BOLD fails to identify the representative ASV sequence of this cluster although *N. roborana* is represented in BOLD by one BIN and 24 barcoded specimens, all from Europe.

**Noctuidae_cluster169:** *Mesoligia furuncula*

This appears to be a numt of *M. furuncula*.

**Tortricidae_cluster249:** *Lathronympha strigana*

This appears to be a numt of *L. strigana*.

**Nymphalidae_cluster25:** *Nymphalis ladakensis*

Matches a chromosome assembly. This appears to be a numt of *A. urticae*. All 6 samples with this cluster also have *A. urticae* reads (Nymphalidae_cluster1).

**Nymphalidae_cluster27:** *Nymphalis ladakensis*

Matches a chromosome assembly. This appears to be another numt of *A. urticae*. All 3 samples with this cluster also have *A. urticae* reads (Nymphalidae_cluster1).

**Noctuidae_cluster171:** *Charanyca ferruginea*

This species is known as *Rusina ferruginea* in DynTaxa. The representative ASV matches a chromosome sequence in GenBank at 99.28% identity. The second closest hit is a CO1 sequence of the same species at 96.88%. The species is well represented in BOLD (27 barcoded specimens in one BIN) but is not represented in IBA except for this strange cluster. BOLD does not identify the sequence but gives *C. ferruginea* as the closest hit with 97.67% similarity. A mystery.

**Ypsolophidae_cluster13:** *Ypsolopha lucella*

This appears to be a numt of *Y. lucella*.

**Nymphalidae_cluster28:** *Nymphalis ladakensis*

Matches a chromosome assembly. This appears to be yet another numt of *Aglais urticae*. All three samples contain CO1 reads of *A. urticae*.

**Nepticulidae_cluster58:** *Zimmermannia liebwerdella*

The match at only 94.70% suggests that this is not *Z. liebwerdella*. The OTU does not co-occur with any of the two other *Zimmermannia* OTUs in the IBA data. It seems possible at least that this represents a new or previously unsequenced species, although the number of reads is small (120 reads in 2 samples). *Z. liebwerdella* splits into three very distinct clusters in BOLD, each of which probably represents a distinct species. Nepticulidae_cluster58 is closest to the North European BIN that is a *Fagus* (beech) feeder. The genus is taxonomically challenging in a broader geographic context. They are also not well studied because they are hard to find as they mine bark. Several other nominal species in the genus, such as *Z. atrifrontella* and *Z. amani* (the other two species encountered in the IBA data) also show deep intraspecific splits in barcode data.

**Tortricidae_cluster262:** *Capua vulgana*

At 96.17% identity, this OTU appears unlikely to represent *C. vulgana*. The latter species seems to be quite variable; it is represented in IBA by six clusters (Tortricidae_cluster246, 288, 291, 292, 295 and 298), all of them matching to the same BOLD BIN with the annotation *C. vulgana*. Therefore, it seems possible at least that this OTU could represent yet another variant. There is no occurrence overlap with any of the samples containing *C. vulgana* OTUs, however. The read and sample numbers are low across the board.

**Nymphalidae_cluster29:** *Nymphalis urticae*

The match is to a nuclear genome assembly. The OTU co-occurs in all three samples with CO1 sequences of *Aglais urticae*. This appears to be yet another numt of *Aglais urticae*.

**Geometridae_cluster187:** *Selenia dentaria*

The match is to a nuclear genome assembly. All four samples also contain CO1 sequences of *S. dentaria*. This appears to be a numt of *S. dentaria*.

**Nepticulidae_cluster64:** *Stigmella anomalella*

This appears to be a Sintax annotation failure.

**Lycaenidae_cluster11:** *Aricia artaxerxes*

This appears to be a Sintax annotation failure.

**Geometridae_cluster189:** *Geometra papilionaria herbacearia*

This appears to be a numt of *Geometra papilionaria*. The single sample containing this OTU also contains a large number of true CO1 sequence reads of *G. papilionaria*.

**Crambidae_cluster54:** *Udea decrepitalis*

At 96.89% identity, this seems unlikely to represent *U. decrepitalis*, nor seems it likely that it would represent *U. inquinatalis*, which is almost as close. This cluster is present in three IBA samples, one of which also contains a cluster that is annotated as *U. decrepitalis*. This could potentially represent a new or unsequenced species, but the possibility that it is a rare variant or numt of *U. decrepitalis* cannot be dismissed.

**Noctuidae_cluster174:** *Mythimna impura*

This appears to be a numt of *M. impura*.

**Erebidae_cluster48:** *Thumatha senex*

This appears to be a numt of *T. senex*.

**Gelechiidae_cluster122:** *Bryotropha terrella*

This appears to be a numt of *B. terrella*.

**Coleophoridae_cluster62:** *Coleophora badiipennella*

At 93.30% identity, this appears unlikely to represent *C. badiipennella*. This species is not present otherwise in IBA. However, cluster 62 occurs in two samples, and in both samples together with reads of other *Coleophora* OTUs. Perhaps this is a "deep" numt in *Coleophora*, or it might represent a new or unsequenced species.

**Tortricidae_cluster271:** *Dichrorampha tarmanni*

At 95.45% identity, this does not appear to be *D. tarmanni*. This species is not represented otherwise in IBA but there is a number of other species in the same genus with similar BLAST scores to *D. tarmanni*. It is possible that this OTU is a numt of one of these, or a deep numt in Dichrorompha. However, it cannot be excluded that this OTU represents a new or unsequenced species.

**Cosmopterigidae_cluster8:** *Sorhagenia rhamniella*

This appears to be a numt of *S. rhamniella*.

**Pterophoridae_cluster23:** *Amblyptilia punctidactyla*

This appears to be a numt of *A. punctidactyla*.

**Noctuidae_cluster181:** *Mythimna conigera*

This appears to be a numt of *M. conigera*.

**Crambidae_cluster56: Elophila nymphaeata**

This appears to be a numt of *E. nymphaeata*.

**Tortricidae_cluster284:** *Epinotia trigonella*

This appears to be a numt of *E. trigonella*.

**Geometridae_cluster200:** *Cyclophora porata*

There are only 13 reads in one sample of this OTU. At 97.85% identity, it might represent *C. porata*, which is a Swedish species that is otherwise not present in the IBA data.

**Noctuidae_cluster186:** *Mythimna straminea*

This appears to be a numt of *M. straminea*.

**Noctuidae_cluster188:** *Xestia sexstrigata*

This appears to be a numt of *X. sexstrigata*.

**Plutellidae_cluster9:** *Ypsolopha sylvella*

At 94.98% identity, this appears unlikely to represent *Y. sylvella*, which has not been recorded in IBA. However, it may well represent a numt of one of the many *Ypsolopha* species encountered in IBA.

**Tortricidae_cluster290:** *Dichrorampha petiverella*

This appears to be a numt of *D. petiverella*.

**Geometridae_cluster202:** *Abraxas grossulariatus*

Potentially, this could represent *A. grossulariatus*, a Swedish species that is not otherwise represented in the IBA data. However, the failed Sintax annotation and the top BLAST hit having only 96.84% identity is strange. The cluster only occurs in one sample, and those samples do not contain the other Swedish species in the genus, *A. sylvanus*, which is present in two IBA samples in low read numbers.

**Nepticulidae_cluster75:** *Stigmella prunetorum*

This seems to be a Sintax annotation failure.

**Elachistidae_cluster74:** *Depressaria incognitella*

This might potentially be a Sintax annotation failure but then this is a new record for Sweden. The distance is also a little far (97.37% identity). This OTU could also be a numt of some other *Depressaria* OTU in IBA.

**Gracillariidae_cluster84:** *Phyllonorycter quercifoliella*

This appears to be a numt of *P. quercifoliella*.

**Noctuidae_cluster190:** *Chortodes pygminus*

This appears to be a numt of *Denticucullus pygmina* (Noctuidae_cluster99). Both occur in only a single sample, together. There are 5 reads of this OTU and 8,720 reads of true CO1 cluster representing *D. pygmina*.

**Gelechiidae_cluster131:** *Athrips pruinosella*

This appears to be a numt of *A. pruinosella*.

**Gracillariidae_cluster89:** *Acrocercops brongniardella*

This appears to be a numt of *A. brongniardella*.

#### Analysis of unclassified clusters

We also analyzed the nochimera swarm clusters (OTUs) of Lepidoptera, which were either not resolved to family ("unclassified.Lepidoptera") or not annotated at the family level ("Lepidoptera_X"). These clusters were filtered away as likely noise in our pipeline but we nevertheless wanted to analyze what these clusters might represent. We used the same procedure for analyzing these clusters as for analyzing the unidentified cleaned Lepidoptera clusters in the previous analysis. All online searches for this analysis were performed on 2023-06-13. Species names associated with clusters represent closest matches in GenBank.

**Lepidoptera_X_cluster1:** BOLD:ACW2340, *Bucculatrix humiliella*

At 97.13% identity, this is fairly unlikely to be *B. humiliella*. The latter species is in Bucculatricidae_cluster12. Lepidoptera_X_cluster1 is a fairly abundant OTU, so unlikely to represent a numt. Only 2 of the 16 samples in which it occurs includes any reads from other *Bucculatrix* species, and those samples include *B. humiliella* but in lower read numbers than this OTU. The BIN it matches to is represented in BOLD by a single undetermined Lepidoptera specimen from Spain. This could represent a new or previously unsequenced species.

**Lepidoptera_X_cluster2:** BOLD:ACB0733, *Scythris potentillella*

At 96.89% identity, this is fairly unlikely to be *S. potentillella*, which is in Scythrididae_cluster4. Lepidoptera_X_cluster2 is a fairly abundant OTU, so unlikely to represent a numt. It does not co-occur with any other OTU placed in *Scythris*. The BIN it matches in BOLD is represented by a single undetermined Lepidoptera specimen from Spain. This could represent a new or previously unsequenced species. *Scythris potentillella* has a very close relative *S. cicadella*. However, the single sequence of this species in BOLD is even more different. It is possible that this represents a cryptic species but it could also be a case of a deep intraspecific split.

**Lepidoptera_X_cluster3:** BOLD:ABV7960, *Coleophora otidipennella*

At 97.13% identity, this is fairly unlikely to be *C. otidipennella*, which seems to be in **Coleophoridae_cluster23** in IBA (see analysis of unidentified clusters). **Lepidoptera_X_cluster3** occurs with 558 reads in one sample, which seems an unlikely large number of reads for a numt. The same sample does contain other *Coleophora* sequences, but no sequences of Coleophoridae_cluster23. The read numbers are fairly equal (558 versus 1,288), which also suggests that this is not a numt. It could be a case of rare local heteroplasmy. However, the BIN it matches is represented in BOLD by a single undetermined Lepidoptera specimen from Poland, so this could potentially also represent a new species.

**Lepidoptera_X_cluster4:** BOLD:ACL9793, *Eupithecia icterata*

This appears to be a numt of *E. icterata*.

**Lepidoptera_X_cluster5:** BOLD:AAI2708, *Platytes cerussella*

At 96.65% identity, this is relatively unlikely to be *P. cerussella*. The latter species is not present in IBA. Lepidoptera_X_cluster5 occurs in two samples (238 and 6 reads, respectively). There are other *Platytes* reads in the first sample, but in approximately equal numbers (380 reads), suggesting that it is not a numt of *Platytes*. The BIN it matches is represented in BOLD by a single undetermined Lepidoptera specimen from Romania. This could potentially represent a new species.

**Lepidoptera_X_cluster6:** BOLD:ABA4188, *Elachista cycotis* (or *E. infuscata*)

At 96.65% identity, this is relatively unlikely to be *E. cycotis*, which is an Australian species. **Lepidoptera_X_cluster6** occurs in 15 samples but with a small number of reads in each (381 reads in total). There are other *Elachista* reads in only five of these samples, suggesting that this is not a numt in *Elachista*. The BIN it matches most closely is represented in BOLD by nine specimens, seven of which are from Italy and identified as *Elachista infuscata* (the remaining two are unidentified). This could possibly represent the latter species, which belongs to the so-called "Cosmiotes" group of *Elachista*. They are very similar in morphology and one species in the group could easily have remained unnoticed. However, *E. infuscata* is a rather distant species occurring in the Mediterranean/Alps area. On the other hand, this species is also known from Siberia (Lauri Kaila, pers. comm.), suggesting that its presence in northern Europe cannot be excluded. The group is taxonomically difficult and the divergences rather small, so a possibility exists that the barcode is shared with some other species in the group, such as *E. freyerella*.

**Lepidoptera_X_cluster7:** BOLD:AAY8772, *Cydia duplicana*

This appears to be a numt of *C. duplicana*.

**Lepidoptera_X_cluster8:** BOLD:ACW8519, *Polix coloradella*

The BIN that this cluster matches to now has three specimens in BOLD, two of which are from Norway and identified as *Denisia albimaculea*, a species that occurs in Sweden.

**Lepidoptera_X_cluster9:** BOLD:ACX1163, *Stigmella myrtillella*

At 92.11% identity, this is unlikely to be *S. myrtillella*. The latter species is not in IBA. **Lepidoptera_X_cluster9** occurs in 2 samples, with a small number of reads in each (18 and 9, respectively). There are other *Stigmella* reads in both samples, not huge numbers but significantly larger numbers (4.2 times and 16 times), suggesting that this could possibly be a numt of *Stigmella*. The genus is rarely represented by a large number of reads, potentially explaining why the numt has gone undetected in most of the *Stigmella* samples (there are more than 1000 of them in total). The BOLD BIN matching this OTU includes only one specimen, an undetermined Lepidoptera specimen from Austria. The possibility that this OTU represents a new or unsequenced species cannot be completely dismissed without closer genetic analysis.

**Lepidoptera_X_cluster10:** BOLD:AAW7369, *Bucculatrix notella*

At 94.72% identity, this is unlikely to be *B. notella*. The latter species is not in IBA. Lepidoptera_X_cluster10 occurs in 11 reads in 1 sample. There are 73 reads of other *Bucculatrix* species in this sample, suggesting that this could possibly be a numt of *Bucculatrix*. The BOLD BIN matching this OTU includes four undetermined Lepidoptera specimens from Estonia, Armenia and Denmark. Without further analysis, it is difficult to determine whether this OTU might represent a new or unsequenced species, or a numt.

**unclassified.Lepidoptera_cluster1:** *Agriphila straminella*

This apparently represents a common CO1 variant in *A. straminella* or a separate species, which is cryptic enough that specimens are typically determined as the same species as *A. straminella*. It occurs in 27 samples, 21 of which also contains the most common cluster of A. straminella CO1 sequences (Crambidae_cluster3, see analysis of BOLD annotation problems). The read ratio range of cluster 3 to this cluster is (0.004, 33.8), so either the frequency of this variant varies a lot, or it represents a separate species. Compare with the complete overlap (5 of 5 samples) and stable ratio range (1522, 9687) of unclassified.Lepidoptera_cluster53, which obviously is a numt of *A. straminella*. The fairly large co-occurrence with *A. straminella* main variant suggests that it might be a case of heteroplasmy.

**unclassified.Lepidoptera_cluster2:** *Agriphila straminella*

This OTU appears to be about 4% different from the previous one, and the case is similar. It occurs in 55 samples, 42 of which also contain the dominant *A. straminella* CO1 cluster (Crambidae_cluster3). The read ratio varies a lot (0.005,1029).

**unclassified.Lepidoptera_cluster3:** *Lathronympha strigana*

At 92.58% identity, this is unlikely to be *L. strigana*. The OTU appears in 5 samples, 4 of which also contain *L. strigana* but in widely varying read number ratios (0.01,71), suggesting that it is not a numt of this species. Widening the taxonomic scope of a possible CO1 parent to the genus *Lathronympha* does not change the overlap analysis. This OTU could potentially represent a new or unsequenced species.

**unclassified.Lepidoptera_cluster4:** *Ancylis myrtillana*

This is apparently a numt of *A. myrtillana*. There are 40 samples containing this OTU and 39 of them contain apparently true CO1 sequences of *A. myrtillana* in much larger numbers (12.4 to 1,515 times more reads). The only sample with cluster 4 that does not contain *A. myrtillana* CO1 sequences contains only 6 reads, so this could potentially be due to contamination.

**unclassified.Lepidoptera_cluster5:** *Sesia apiformis*

Matches a nuclear genome assembly at 100.00% identity. This is apparently a numt of *S. apiformis*.

**unclassified.Lepidoptera_cluster6:** *Stigmella prunetorum*

This seems to be a Sintax annotation failure. This cluster is very close to Nepticulidae_cluster75, which can be assigned with some confidence to *S. prunetorum*. The current cluster is actually identical to GenBank records (100.00% identity), while cluster 75 only has about 98.6% identity to the same entries. The swarm clustering procedure would presumably have grouped these ASVs in the same cluster if it were not for the fact that the family annotation problem resulted in swarm clustering being run separately for the two ASV sets.

**unclassified.Lepidoptera_cluster7:** *Eucosma aspidiscana*

This appears to be a Sintax annotation failure. The representative ASV sequence is close to Tortricidae_cluster179, and Tortricidae_cluster210 both of which appear to belong to *E. aspidiscana* or possibly a close relative (see analysis of unidentified clusters). There is no occurrence overlap between these three clusters.

**unclassified.Lepidoptera_cluster8:** *Bucculatrix cristatella*

At 97.12% identity, this may be different from *B. cristatella*. This OTU occurs in 1 sample with 2561 reads, while the main form of *B. cristatella* is in two IBA clusters, Bucculatricidae_cluster1 and Bucculatricidae_cluster2. The ASVs are assigned to a unique BOLD BIN in the first of these, and to two different BOLD BINs in the second (including a few ASVs that apparently cannot be assigned to either of these two). The annotation failure for unclassified.Lepidoptera_cluster8 indicates that it has some feature linking it to taxa outside of Bucculatricidae. The most likely interpretation appears to be that this is a CO1 variant of *B. cristatella*, but it could also represent a cryptic species.

**unclassified.Lepidoptera_cluster9:** *Stigmella confusella*

This appears to be a Sintax annotation failure (the species epithet appears appropriate!). The species is recorded from Sweden.

**unclassified.Lepidoptera_cluster10:** *Lycaena virgaureae*

This appears to be a Sintax annotation failure. The species occurs in Sweden.

**unclassified.Lepidoptera_cluster11:** *Ypsolopha parenthesella*

This is apparently a numt of *Y. parenthesella*.

**unclassified.Lepidoptera_cluster12:** *Charissa obscurata*

This appears to be a Sintax annotation failure. The species occurs in Sweden.

**unclassified.Lepidoptera_cluster13:** *Stigmella lapponica*

This appears to be a Sintax annotation failure. The species occurs in Sweden.

**unclassified.Lepidoptera_cluster14:** *Trifurcula cryptella*

This could be a Sintax annotation failure, even though the top hit score is low at 97.37%. Alternatively, it could be another species in the genus. The OTU occurs in 6 samples with no overlap with the other *Trifurcula* OTU (*Trifurcula immundela*, Nepticulidae_cluster70) in the IBA darta (see also unclassified.Lepidoptera_cluster56 below). The species occurs in Sweden.

**unclassified.Lepidoptera_cluster15:** *Archips betulanus*

This could be a Sintax annotation failure, even though the top hit score is low at 96.89%. The species occurs in Sweden.

**unclassified.Lepidoptera_cluster16:** *Stigmella anomalella*

This appears to be a Sintax annotation failure. The species occurs in Sweden.

**unclassified.Lepidoptera_cluster17:** *Philedonides lunana*

This appears to be a Sintax annotation failure for a slightly divergent *P. lunana* ASV sequence, probably represented by a single specimen. The species appears in several IBA clusters and BOLD BINs.

**unclassified.Lepidoptera_cluster18:** near *Lampronia aenescens*

At 93.06% identity, this cluster is unlikely to be *L. aenescens*. There are 448 reads of this OTU in one sample; the same sample contains 6 reads of *L. redimitella*. This seems to potentially represent a new or unsequenced species of *Lampronia*.

**unclassified.Lepidoptera_cluster19:** near *Parornix traugotti*

This could be a Sintax annotation failure, even though the top hit score is low at 97.37%, especially given that it has been sequenced from Denmark and Lithuania. This OTU occurs in 3 samples, 2 of which also contain reads of other *Parornix* OTUs. The ratio between read numbers is somewhat equal. Both the partial overlap and the read ratios suggest that this is not a numt. This could potentially represent a new species close to *P. traugotti*, which occurs in Sweden. It could also be a previously unknown haplotype of *P. traugotti*, or even *P. polygrammella*, which is split into two BINs in BOLD (both BINs are grouped in the same OTU cluster in our pipeline).

**unclassified.Lepidoptera_cluster20:** near *Stigmella pretiosa*

This appears to be a Sintax annotation failure. The species occurs in Sweden.

**unclassified.Lepidoptera_cluster21:** near *Dichrorampha simpliciana*

This is apparently a numt of *D. simpliciana*.

**unclassified.Lepidoptera_cluster22:** near *Nymphalis ladakensis*

This is apparently yet another numt of *Aglais urticae*. The top hit is in a nuclear genome assembly and the OTU is present in 4 samples, in all cases together with large numbers of *A. urticae* CO1 sequences.

**unclassified.Lepidoptera_cluster23:** near *Recurvaria leucatella*

This is apparently a numt of *R. leucatella*.

**unclassified.Lepidoptera_cluster24:** near *Elachista freyerella*

At 97.61% identity, this is not so likely to be *E. freyerella*, although the latter is a Swedish species that is otherwise absent in the IBA data. The OTU has only a total of 148 reads in 8 samples, suggesting that it might be a numt. All 8 samples contain a large number of reads of Elachistidae_cluster1, which appears to be *E. exactella* (see analysis of unidentified clusters). The read ratio range is also suggestive of a numt (9.7, 21493).

**unclassified.Lepidoptera_cluster25:** near *Orthosia cruda*

This is apparently a numt of *O. cruda*.

**unclassified.Lepidoptera_cluster26:** near *Ypsolopha mucronella*

This is apparently a numt of *Y. mucronella*.

**unclassified.Lepidoptera_cluster27:** near *Choristoneura hebenstreitella*

This is apparently a numt of *C. hebenstreitella*.

**unclassified.Lepidoptera_cluster28:** near *Ochsenheimeria urella*

At 91.15% identity, this appears unlikely to be *O. urella*. The OTU occurs in 2 samples with a total of 79 reads. There is no overlap with any other OTU annotated as belonging to the genus *Ochsenheimeria*. This might potentially represent a new or unsequenced species.

**unclassified.Lepidoptera_cluster29:** near *Chilodes maritimus*

This could be a numt of *Chilodes maritimus* (Noctuidae_cluster138) or perhaps a CO1 variant of this species (98.33% identity to a claimed mitochondrial sequence of *C. maritimus* in GenBank). Note that the name form varies, so that it is given as *C. maritima* (probably incorrectly) in DynTaxa and the Sintax annotation. The two OTUs occur in one and the same sample. This OTU has 129 reads, the other 2234 reads. There is a single ASV in each OTU.

**unclassified.Lepidoptera_cluster30:** near *Ypsolopha mucronella*

This is apparently a numt of *Y. mucronella*.

**unclassified.Lepidoptera_cluster31:** near *Acrobasis marmorea*

This is apparently a numt of *A. marmorea*.

**unclassified.Lepidoptera_cluster32:** near *Agriphila inquinatella*

This is apparently a numt of *A. inquinatella*.

**unclassified.Lepidoptera_cluster33:** near *Mythimna impura*

The top hit is in a nuclear genome assembly. This is apparently a numt of *M. impura*.

**unclassified.Lepidoptera_cluster34:** near *Ypsolopha mucronella*

This is apparently a numt of *Y. mucronella*.

**unclassified.Lepidoptera_cluster35:** near *Chionodes lugubrella*

This is apparently a numt of *C. lugubrella*.

**unclassified.Lepidoptera_cluster36:** near *Pseudotelphusa scalella*

This is apparently a numt of *P. scalella*.

**unclassified.Lepidoptera_cluster37:** near *Lypusa maurella*

This appears to be a numt of *L. maurella* (in Lypusidae_cluster5). It occurs in 3 samples together with the latter, read number ratio range (30.3,65.8).

**unclassified.Lepidoptera_cluster38:** near *Ypsolopha mucronella*

This is apparently a numt of *Y. mucronella*.

**unclassified.Lepidoptera_cluster39:** near *Epirrita christyi*

This could be a numt of *E. christyi* but could also be a rare variant. It occurs in 3 samples, 2 of which also contain *E. christyi* reads at a dominance ratio range (12.4,12.7). The two samples where the two OTUs co-occur have a relatively low number of reads of *E. christyi* compared to the other ones, perhaps suggesting that the specimens in those samples are heteroplasmic.

**unclassified.Lepidoptera_cluster40:** near *Ypsolopha mucronella*

This is apparently a numt of *Y. mucronella*.

**unclassified.Lepidoptera_cluster41:** near *Gypsonoma adjuncta*

At 92.33% identity, this appears unlikely to represent *G. adjuncta*. The OTU occurs in 2 samples with a total of 44 reads. The same samples contain a range of (54.7, 137.6) times as many sequences of other *Gypsonoma* OTUs, suggesting that this might be a *Gypsonoma* numt.

**unclassified.Lepidoptera_cluster42:** near *Batia lunaris*

This is apparently a numt of *B. lunaris*.

**unclassified.Lepidoptera_cluster43:** near *Sophronia sicariellus*

This is apparently a numt of *S. sicariellus*.

**unclassified.Lepidoptera_cluster44:** near *Micropterix sp. BOLD:AAI1534*

At 93.54% identity, this is unlikely to be a match to BOLD:AAI1534. This OTU could be a numt of some *Micropterix* species. There are 19 reads of the OTU in one sample. The same sample has 2,751 reads of other OTUs assigned to *Micropterix*.

**unclassified.Lepidoptera_cluster45:** near *Batia lunaris*

This is apparently a numt of *B. lunaris*.

**unclassified.Lepidoptera_cluster46:** near *Batia lunaris*

This is apparently a numt of *B. lunaris*.

**unclassified.Lepidoptera_cluster47:** near *Lathronympha strigana*

This is apparently a numt of *L. strigana*.

**unclassified.Lepidoptera_cluster48:** near *Ypsolopha mucronella*

This is apparently a numt of *Y. mucronella*.

**unclassified.Lepidoptera_cluster49:** near *Batia lunaris*

This is apparently a numt of *B. lunaris*.

**unclassified.Lepidoptera_cluster50:** near *Pleurota bicostella*

This is apparently a numt of *P. bicostella*.

**unclassified.Lepidoptera_cluster51:** near *Ypsolopha mucronella*

This is apparently a numt of *Y. mucronella*.

**unclassified.Lepidoptera_cluster52:** near *Ypsolopha mucronella*

This is apparently a numt of *Y. mucronella*.

**unclassified.Lepidoptera_cluster53:** near *Agriphila straminella*

This is a numt of *A. straminella*. The top hit is in a nuclear genome assembly, and the OTU is present in small numbers in 4 samples that all contain very large numbers of true CO1 sequences of the same species (Crambidae_cluster3, see analysis of BOLD annotation problems).

**unclassified.Lepidoptera_cluster54:** near *Micropterix tunbergella*

This is apparently a numt of *M. tunbergella*.

**unclassified.Lepidoptera_cluster55:** near *Ypsolopha parenthesella*

This is apparently a numt of *Y. parenthesella*.

**unclassified.Lepidoptera_cluster56:** near *Trifurcula cryptella*

At 94.02% identity, this appears unlikely to be *T. cryptella*. Instead it appears to be a numt of unclassified.Lepidoptera_cluster14 (potentially *Trifurcula cryptella*), see above. Cluster 56 occurs in only one sample; the same sample has 51 times more reads of cluster 14 (308 versus 6 reads). The same sample is also close to the top read number for cluster 14 (308 versus 337 reads).

**unclassified.Lepidoptera_cluster57:** near *Ypsolopha rhinolophi*

At 95.22% identity, this appears relatively unlikely to represent *Y. rhinolophi*, which does not occur in Sweden (it is recently described from northern Portugal and south-east France). Therre are only 5 reads of this OTU in one sample. In the same sample, there are 2,672 reads of other *Ypsolopha* OTUs, suggesting that the current OTU may be a numt of some *Ypsolopha* species.

**unclassified.Lepidoptera_cluster58:** near *Catoptria falsella*

This is apparently a numt of *C. falsella*.

**unclassified.Lepidoptera_cluster59:** near *Agriopis marginaria*

This is apparently a numt of *A. marginaria*.

#### Analysis of range expansions

[Either include Rmd file content here, or add all of this document to the Rmd file, or just insert a reference here to the Rmd file, ideally with a short summary of this analysis.]

### Supplementary Results S1

In the following, we discuss the detailed results for the clusters selected for validation (marked in yellow above). The list includes OTU clusters identified as species that are new to Sweden or potentially new to science, as well as OTU occurrence records that are well outside the known range of the corresponding species in Sweden. The analysis underlying the selection of each species is given as follows: (#1) Analysis of Dyntaxa misses; (#2) Analysis of BOLD annotation problems; (#3) Analysis of unidentified clusters; (#4) Analysis of unclassified clusters; and (#5) Analysis of range expansions. Similarity values are from online blastn searches against GenBank.

#### Species potentially new to Sweden

**Geometridae_cluster181**: *Eupithecia groenblomi* (#1)

We investigated the single sample - SAWGVX - that contained this cluster; in total there were 216 reads of it in the sample. Three specimens of *Eupithecia*, which could represent *E. groenblomi*, were selected for barcoding, and all of them yielded full-length barcodes. Two of them matched *Eupithecia tantillaria*, a species expected in the sample because it has many reads (7,691) for Geometridae_cluster48, annotated by our pipeline as *E. tantillaria*. The third specimen matched *E. groenblomi* with 100% similarity. The results therefore confirm that this previously unrecorded, difficult to find species is present in Sweden.

**Geometridae_cluster186**: *Agriopis budashkini* (#1)

We analyzed sample SNDQSZ, which contained 90 reads of Geometridae_cluster186. There were no adults that could correspond to *A. budashkini* but there was a small geometrid larva, which could potentially belong to the species. The full barcode of the larva had 99% similarity to *A. budashkini*, suggesting that the identification is correct.

**Gracillariidae_cluster61**: *Phyllonorycter mespilella* (#1)

We sorted the single sample - SCUQKH - containing this cluster. The full-length barcode of the specimen was 100% identical to *P. mespilella*.

**Batrachedridae_cluster3**: *Batrachedra confusella* (#1)

We selected one sample for validation – S9IQKE. It had a good number of Batrachedridae_cluster3 reads (539); no other Batrachedridae cluster was present. Two potential specimens were selected for barcoding but both turned out to be *Coleophora glaucicolella*, which was expected in the sample based on the presence of Coleophoridae_cluster4, annotated as *C. glaucicolella*, with 289 reads. Thus, we did not find *B. confusella* specimens in this sample but it is still possible that it is present in other samples.

#### Species potentially new to science

**Lepidoptera_X_cluster1**: BOLD:ACW2340, near *Bucculatrix humiliella* (#3)

We sorted one sample (SFD7FT) and found a number of specimens of *Bucculatrix* (10 individuals). The full-length barcodes of these specimens were ~3% different from known barcodes of *B. humiliella.* This suggests that this could indeed be a new *Bucculatrix* species, although it could also represent a new and previously unrecorded CO1 variant of *B. humiliella*.

**Lepidoptera_X_cluster2:** BOLD:ACB0733, near *Scythris potentillella* (#3)

We selected one sample (S9TT4P) for verification; it had 188 reads for this cluster. Sorting revealed two potential individuals matching this species. Both specimens were successfully barcoded and had the same sequence, 97% similar to *S. potentillella.* This confirms the presence of two individuals in this sample, representing a CO1 variant that has not been included in BOLD yet. Potentially, this could represent a new species.

**Lepidoptera_X_cluster5:** BOLD:AAI2708, near *Platytes cerussella* (#3)

We looked at one sample (SBCM1Z) with 238 reads and analyzed the three potential candidate specimens in this sample. Barcoding showed that they were all *Platytes cerusella* (99% to 100% similarity), a species that was expected from the sample due to the presence of Crambidae_cluster40 (380 reads), annotated to *P. cerusella*. Thus, it seems likely that Lepidoptera_X_cluster5 represents a numt or heteroplasmic variant of CO1, which did not amplify using the primers for the full Folmer region.

**Lepidoptera_X_cluster6:** near BOLD:ABA4188, near *Elachista cycotis* (or *E. infuscata*) (#3)

One sample was sorted (SMTMDU), and we found one potential specimen that could represent this species. However, barcoding revealed that it was *Depressaria olerella*, which was expected in this sample based on the presence of Elachistidae_cluster42 (1,141 reads). A potential explanation is that the current cluster represents a numt in the Elachistidae that occurs in a larger taxonomic group or outside of the genus *Elachista*, which it matched most closely. The small number of reads (50) could also indicate that there are only DNA traces of a species in this sample, and no specimen.

**unclassified.Lepidoptera_cluster18:** near *Lampronia aenescens* (#3)

We analyzed the single sample containing this cluster (SNJZZC). The sample generated 448 reads of this cluster and 6 (!) reads of Incurvariidae_cluster19, which is assigned to *Lampronia redimitella*. Detailed analysis of the ASV corresponding to Incurvariidae_cluster19 showed that it is quite similar to barcode sequences submitted to BOLD from Norway and Finland of *L.*

*redimitella*. It is unclear why this ASV did not match to *L. redimitella* in the BOLD identification engine, BLAST searches on GenBank, or in our taxonomic annotation pipeline.

After sorting we had only one candidate specimen. The full-length barcode identified it as *Elachista humilis*, presumably corresponding to Elachistidae_cluster17, which is annotated as *E. humilis* and present in the sample. The apparent *Lampronia* sequences in the sample may represent errors, contaminations, or potentially DNA traces of specimens that are not in the sample (even though the large number of reads suggests otherwise).

**unclassified.Lepidoptera_cluster19:** near *Parornix traugotti* (#3)

We chose sample SXLRKG to look into this cluster. The sample contained no other *Parornix* OTU and had 106 reads of this cluster. One specimen was selected as a possible member of *Parornix*. The full-length barcode BLASTed at 97% identity to *Parornix traugotti* (closest match). There are two Gracillariidae OTUs also present in the sample but both are assigned to *Phyllonorycter (P. ulmifoliella and P. anderidae*). Thus, this cluster seems likely to represent a new species close to *P. traugotti*, or a new CO1 variant of this species.

**Gracillariidae_cluster1:** near *Acrocercops tacita* (#4)

To investigate this cluster we sorted two samples. The first sample, SMIYF7, generated 845 reads of this cluster and 26 reads of Gracillariidae_cluster2, *Acrocercops brongniardella* (= *A. brongniardellus*). We identified two specimens of *Acrocercops*; full-length barcoding identified one specimen as *A. brongniardella* and the other as a separate lineage with *A. brongniardella* as the closest match at 96% identity. The second sample, SNJZZC, generated 643 reads of Gracillariidae_cluster1 and none of Gracillariidae_cluster2. Full-length barcoding identified this specimen, again, as a lineage matching at 96% identity to *A. brongniardella*.

After our initial analysis was completed, a new species of *Acrocercops* was described, *A. andreneli* (Nel et al. 2023). There are now some BOLD reference sequences matching our Gracillariidae_cluster1, which are annotated as belonging to this species even though they are mixed in a clade with other sequences identified as *A. brongniardella*. Nevertheless, it appears likely that this cluster represents *A. andreneli*. The species is now recorded from several provinces in Sweden, although there is considerable uncertainty about the distribution of this species and *A. brongniardella* because they were regarded as a single species in the past (Bengtsson 2024). There is some evidence, however, to suggest that *A. andreneli* is an invasive species that spread northwards in Europe and reached Sweden, where it spread northwards from approximately 2017 and onwards (Bengtsson 2024). The IBA data show that *A. andreneli* was well established, and both this species and *A. brongniardella* were widely distributed in Sweden in 2019. Our data represent the most complete picture we have so far of their distributions in the country.

**Adelidae_cluster11**: near *Cauchas fibulella* (#4)

We looked at the sample SCVLQL with 410 reads of this cluster, and found three specimens that could match. The full-length barcodes all BLAST to *C. fibulella* at 96–97% identity. There were two SNPs between specimens, indicating that there is some genetic variation within the cluster. Nevertheless, the results confirm that this is not a NUMT and suggest that it could be a new species close to *C. fibulella*. An alternative possibility is that the frequently barcoded *C. fibulella* has a genetically distinct CO1 variant in Sweden.

**Bucculatricidae_cluster9**: near *Bucculatrix notella* (#4)

We selected one sample (SQ29ID) with 265 reads of Bucculatricidae_cluster9. Two individuals were selected as plausible members of *Bucculatrix*. For both individuals, the best BLAST hit of the full-length barcode was *B. cristatella* at only 94% similarity. There were no other Bucculatricidae OTUs in this sample. This suggests that the cluster represents a new cryptic species.

**Coleophoridae_cluster47**: near *Coleophora otidipennella* (#4)

We selected one sample (SQHWKT), which had 385 reads for this cluster and also three other Coleophoridae clusters present: Coleophoridae_cluster1 (*C. glitzella*, 2,005 reads), Coleophoridae_cluster4 (*C. glaucicolella*, 847 reads) and Coleophoridae_cluster10 (*C. lusciniaepennella*, 964 reads). Seven individuals, which could potentially represent these clusters, were selected for barcoding. Four matched *Coleophora glitzella*. One matched *Aproaerema/Syncopacma cinctella*, which corresponded to a cluster from a different family, Gelechiidae_cluster61 (161 reads). Another matched *Hypenodes humidalis*, corresponding to Erebidae_cluster7 (1,361 reads). Finally, one specimen (341_G4) yielded a barcode that BLASTed at only 96% identity to *Coleophora otidipennella*. This corroborates that this is a correct CO1 sequence, which could represent a new species or a distinct and previously unknown CO1 variant of *C. otidipennella*.

**Tortricidae_cluster210**: near *Eucosma aspidiscana* (#4)

We selected sample SB7MP8 to look into this cluster; it had 1,570 reads of this cluster. Only one specimen was found as a potential member of *Eucosma*. Barcoding revealed that this was *Ancylis myrtillana*, which corresponds to another OTU in the sample, Tortricidae_cluster2 (545 reads). The large number of reads of Tortricidae_cluster210 might possibly suggest that this is a NUMT or CO1 variant picked up by the metabarcoding primers but not by the full-length primers. However, the correct interpretation remains unclear.

**Nepticulidae_cluster58**: near *Zimmermannia liebwerdella* (#4)

We sorted one sample (S64XAX), which had only 63 reads for this cluster. We found one individual likely to represent this cluster. The barcode of this specimen is 96.8% similar to *Zimmermannia liebwerdella* according to the BOLD identification engine. This confirms the existence of a potential cryptic species or genetically distinct CO1 variant of *Zimmermannia*.

#### Potential range expansions within Sweden

**Coleophoridae_cluster27:** *Coleophora juncicolella*

We checked one sample S5Y7PP (trap_ID: TC8QBB; 66°35'50.4"N 19°50'33.4"E) that had 66 reads for this cluster. Two potential individuals were found. Full-length barcoding showed that one specimen was *C. ledi,* which was expected based on Coleophoridae_cluster16 (644 reads) annotated to this species. The other specimen matched *C. juncicolella*, confirming that this species is present much further north than previously known.

**Coleophoridae_cluster7:** *Coleophora peribenanderi*

We checked one sample – S8UMBT – that had 232 reads for this cluster. One potential individual was found and it was identified with barcoding to be *C. paripennella.* This represents a clustering error in our pipeline, which groups the two species in the same OTU cluster. Most of the reads match *C. peribenanderi*, so this is the resolved name for the cluster. In this particular sample, however, the reads match *C. paripennella*, a closely related but more northern species.

**Geometridae_cluster22**: *Philerema vetulata*

This species was supposed to be in sample SCAM1W but was not found (only 56 reads). This could potentially represent DNA traces of the species, or an error.

**Tortricidae_cluster68**: *Aleimma loeflingiana*

We analyzed two individuals that could represent this cluster from one sample with a large number of reads (10,757) – SPCUGJ. One specimen turned out to be a different species (*Grypsonoma nitidulana*), presumably matching Tortricidae_cluster76 (>7,000 reads) annotated to this species. The other specimen was confirmed to be *Aleimma loeflingiana* with 99% identity.

### Supplementary figures

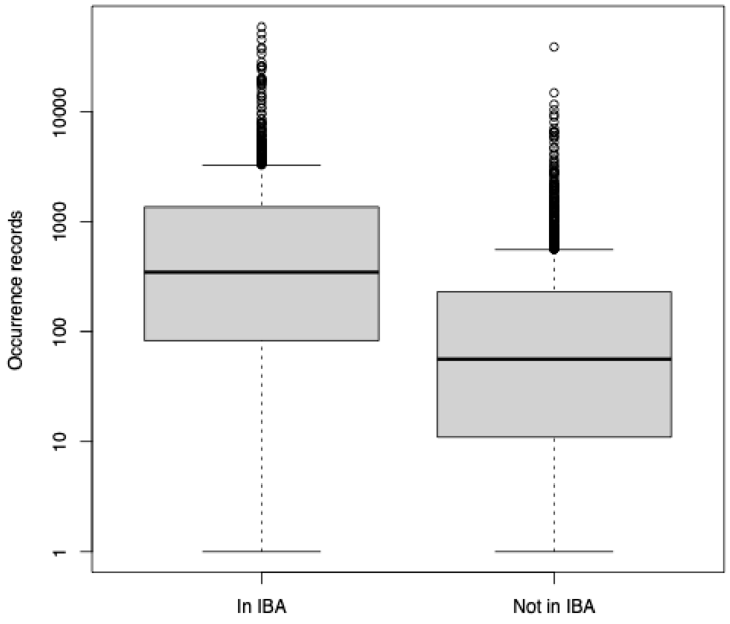


**Fig. S1. Distribution of the total number of GBIF occurrence records for Lepidoptera species detected versus missing in the high-throughput IBA survey.** The species detected in the IBA survey are more frequently reported by naturalists than those that are not recovered.

Even though the variance is large, the difference in mean is significant (log values, two-sided t-test, $p < 2.2e-16$).

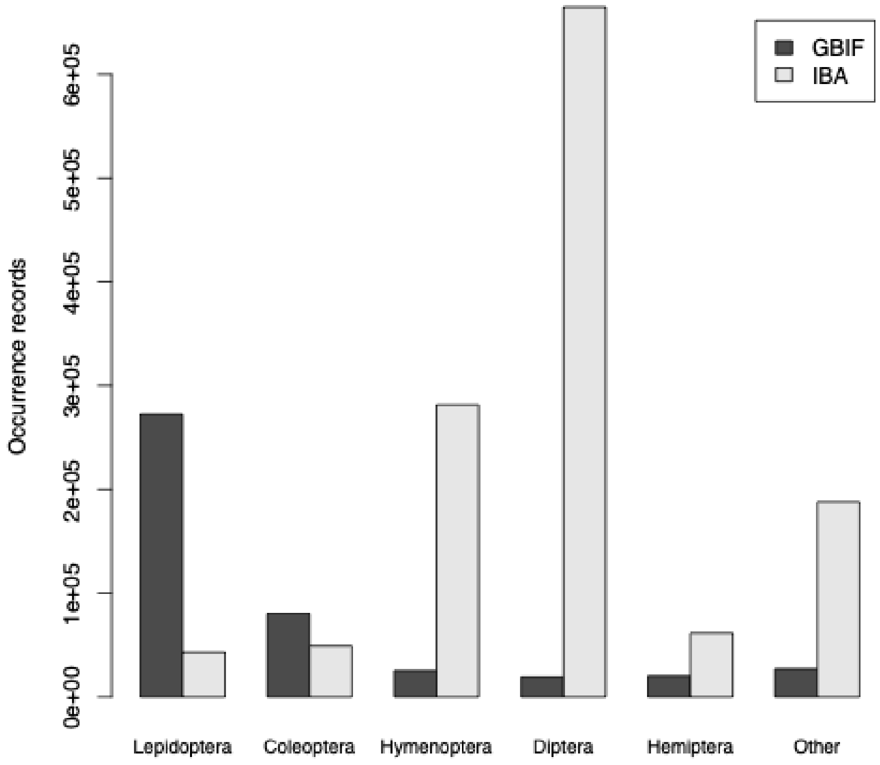


**Fig. S2. Number of occurrence records reported by naturalists in 2019 (GBIF) compared to the high-throughput survey data, viewed as simple occurrence data (IBA).** Among the major insect groups, naturalists only reported substantially more data than generated in the high-throughput survey for Lepidoptera. The naturalists reported slightly more data than IBA for Coleoptera, but only a small fraction of the IBA data for the remaining groups.

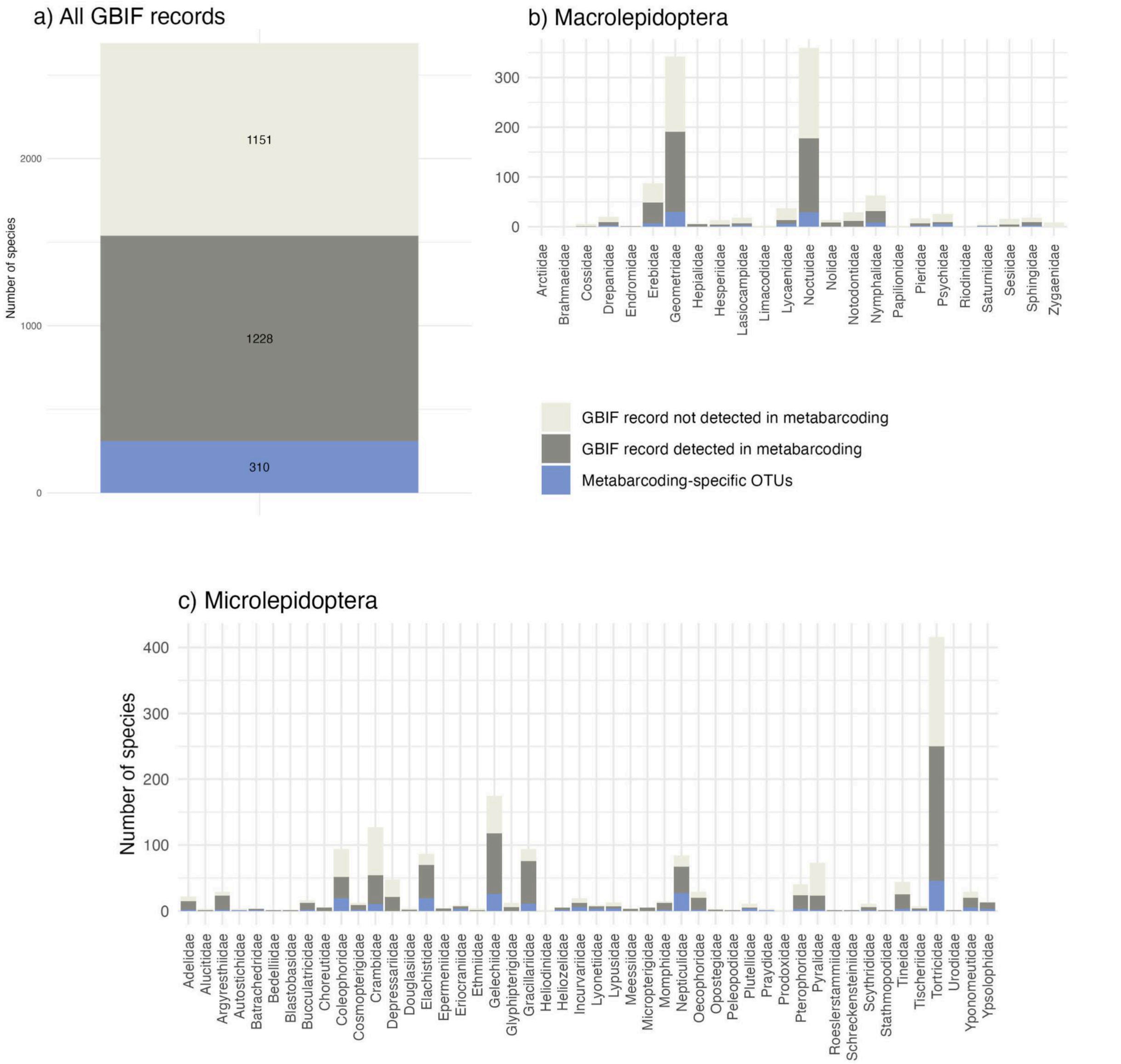


**Fig. S3. The proportion of the species that were reported from Sweden in 2019 by all GBIF data publishers detected by high-throughput methods** A) across all taxa, B) per macrolepidoptera family and C) per microlepidoptera family. Here, the dark gray sections of the bars correspond to Swedish taxa detected by Malaise traps coupled with metabarcoding; the light gray sections show taxa recorded from Sweden in 2019 but not detected by high-throughput methods; and the blue sections show taxa detected uniquely by high-throughput methods. Families are sorted by the traditional (but phylogenetically unsupported) split into “microlepidoptera” and “macrolepidoptera”.

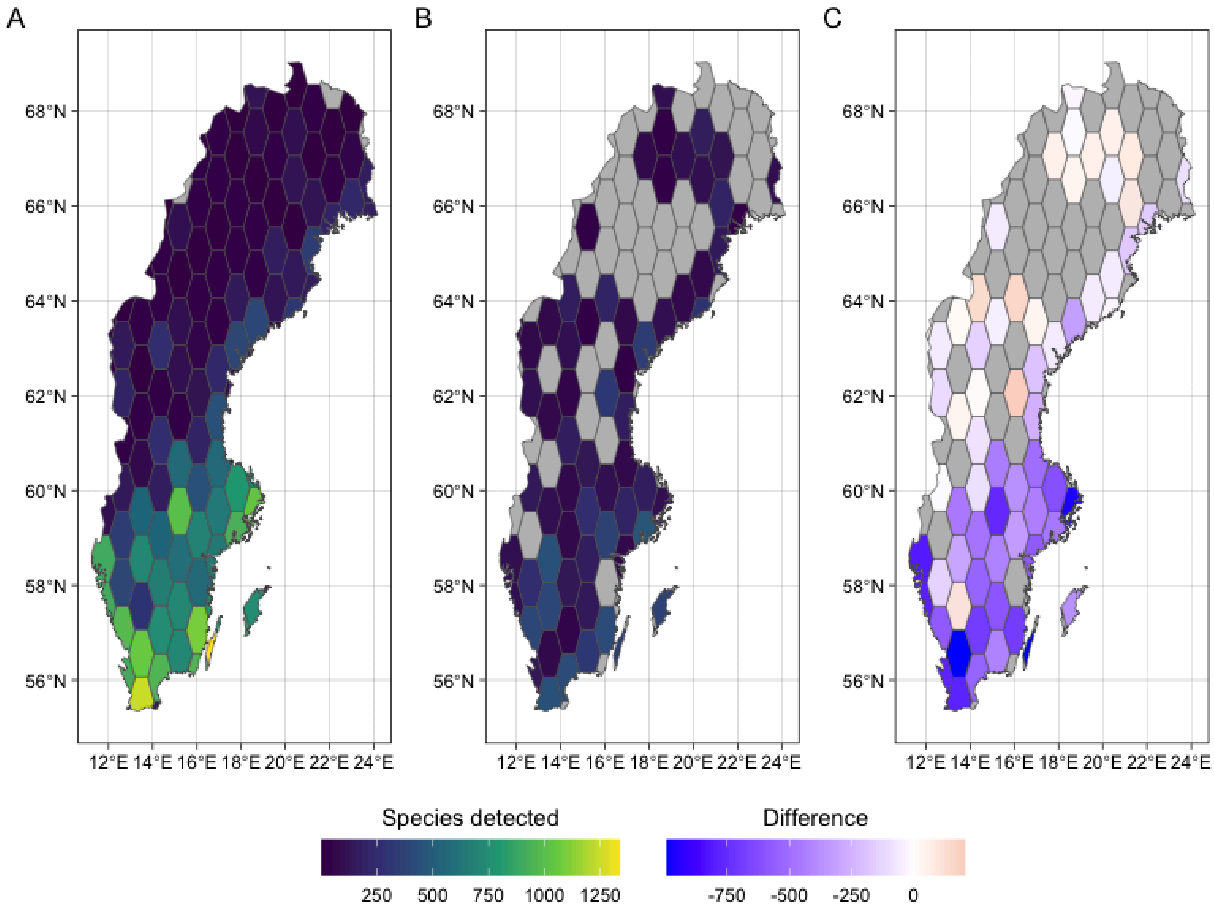


**Fig. S4. Number of species detected in Sweden in 2019.** Numbers are summed across the year and calculated per polygon. Panel (A) shows data from all GBIF data contributors in 2019. Panel (B) summarizes the number of species detected by Malaise trapping coupled with metabarcoding and (C) is the difference between the two (when data for both was available).

### Tables

**Table S1. Numbers of species/OTUs detected and not detected by metabarcoding divided into Macro-, Micro-lepidoptera and butterflies categories.** The known fauna of Sweden excludes *Metabarcoding-specific OTUs*, and therefore is 1227 and 1763 species per Macro- and Micro-lepidoptera respectively. Note that *Butterflies* is a subset of *Macrolepidoptera*.

| | **Macrolepidoptera** | **Microlepidoptera** | **Butterflies (subset of Macrolepidoptera)** |
|---|---|---|---|
| Metabarcoding-specific OTUs | 29 | 45 | 1 |
| Species not detected in metabarcoding | 685 | 770 | 24 |
| Species detected in metabarcoding | 542 | 993 | 8 |
| Percent detected known species [%] | 44.17 | 56.32 | 25 |
